# Intrinsically disordered bHLH family member TCF4 drives LLPS in a DNA-dependent manner

**DOI:** 10.1101/2025.06.18.660356

**Authors:** Nikola Sozańska, Barbara P. Klepka, Anna Niedzwiecka, Beata Greb-Markiewicz, Andrzej Ożyhar, Aneta Tarczewska

**Author notes:** Corresponding author correspondence to Aneta Tarczewska;.

## Abstract

Transcription factor 4 (TCF4), a basic helix-loop-helix (bHLH) protein, plays a critical role in neurodevelopment and is genetically linked to disorders such as Pitt-Hopkins syndrome (PTHS) and schizophrenia. While TCF4 is known to function through sequence-specific DNA binding, the influence of its intrinsically disordered regions (IDRs) on its molecular behavior remains scattered. The TCF4 deletion mutant, lacking a fragment of the bHLH domain, forms puncta in the nucleus, which are reminiscent of patterns observed in disease-associated TCF4 mutants. Our in vitro analyses have demonstrated that the shortest human isoform of TCF4 (TCF4-I^−^), which is predominant in the brain and contains extensive IDRs, undergoes liquid-liquid phase separation (LLPS) *in vitro*. LLPS is promoted by increased ionic strength and molecular crowding, resulting in dynamic, liquid-like condensates that coalesce and display rapid fluorescence recovery after photobleaching (FRAP). Over time, these condensates mature, exhibiting increased viscosity while retaining molecular mobility. Importantly, we show that both specific and non-specific DNA sequences dissolve TCF4 condensates. This suggests that DNA binding competitively inhibits the multivalent interactions necessary for phase separation. These findings indicate that LLPS driven by TCF4 is a reversible process, likely involved in transcriptional activation bursts. We propose that mutations that disrupt DNA binding or localization may trap TCF4 in aberrant condensates, contributing to PTHS pathogenesis through dysregulated phase behavior. Our findings offer a mechanistic link between TCF4 structural features, phase separation, and its transcriptional function, providing novel insight into how mutations in this TF may drive neurological disease through aberrant biomolecular condensation.

## 3. Introduction

It has long been wondered how molecules find their way in a crowded molecular environment so that biochemical reactions can take place. Recent advancements in molecular biology have unveiled that many of them occur in macromolecular condensates, formed due to the capacity of certain protein sequences to form mesoscopic, dynamic, nonstoichiometric supramolecular liquid assemblies [1]. These condensates arise from non-covalent interactions, encompassing Coulomb, hydrophobic, hydrogen bonding, dipole–dipole, π–π, and cation−π interactions among associative protein and nucleic acid molecules [2,3]. The presence of intrinsically disordered regions (IDRs) allows the formation of many weak and transient inter- and intra-chain interactions, essential for the formation of liquid condensates through liquid-liquid phase separation (LLPS) [4–9], the process extensively studied in polymer chemistry, and recently emerged as a notable occurrence in biological systems [3]. LLPS in biology has been shown to be responsible for the creation of more intricate, transient multicomponent assemblies, known as membraneless organelles (MLOs) [1,10,11]. Biophysical studies of numerous MLOs have unveiled their liquid-like nature [12–15]. MLOs exhibit a barrier-less nature, forming spontaneously in response to physicochemical changes in the microenvironment [3] and facilitating the unrestricted exchange of components with the surrounding milieu [1,11].

The importance of the LLPS process in biological systems is described in several recent reports. A growing number of articles indicate that it is a process linked to the proper regulation of gene expression. Precise regulation of transcription is crucial to ensure a rapid cellular response to various internal and external signals. This intricate process relies on the coordinated formation of adaptable complexes involving numerous transcription regulators (TRs), including transcription factors (TFs), cofactors, and chromatin structure regulators. To initiate transcription, TRs must intricately interact within the crowded nuclear environment, precisely aligning with specific DNA sequences. Many eukaryotic TRs contain IDRs, and its remarkable flexibility enables one-to-many interactions that promote the formation of supramolecular complexes [16,17]. It has been shown that short-lived interactions between TFs engage their IDRs, leading to the formation of transient local regions with high concentrations of TFs [18]. In addition, experimental data have provided evidence that some of the Mediator complex subunits and disordered regions of Oct4 (octamer-binding transcription factor 4) and BRD4 (bromodomain-containing protein 4) can form MLOs [19,20]. It has been proposed that LLPS induced by binding of TFs to super enhancers along with Mediator subunits is the mechanism by which cell-specific gene transcription is regulated in vivo [20]. Further evidence suggests that LLPS plays an important role in transcriptional progression and RNA processing [19,21]. FUS (Fused in sarcoma), EWS (Ewing sarcoma) and TAF15 (TATA-binding protein-associated factor 2N) proteins which can form phase-separated, liquid-like condensates, directly bind the C-terminal domain of RNA Polymerase II and activate transcription [22,23]. The C-terminal domain of the RPB1 subunit of human RNA polymerase II consists of a highly repetitive, unstructured low complexity domain (LCD) [24]. In addition, the TRs YAP (yes-associated protein) [25] and TAZ (transcriptional co-activator with PDZ-binding motif) [26] undergo LLPS, which leads to the secretion of transcriptional coactivators and cofactors from the cellular environment, ultimately facilitating the expression of target genes [27].

Furthermore, the LLPS of chromatin-associated factors can promote the organization of the chromatin structure to regulate transcription [28–30]. It appears that reconstructed chromatin undergoes LLPS both *in vitro* and in living cells [31], and heterochromatin protein 1 (HP1) forms condensates through LLPS [28,32]. Some studies indicate that LLPS occurs specifically in transcriptionally active regions of DNA with lower chromatin density. Phase-separated liquid condensates form preferentially in low-density chromatin regions. Thus, distant target genomic *loci* can be pulled toward each other mechanically and restructured by liquid condensate fusion, while other chromatin regions are pushed out mechanically by the condensates [29]. To date, the link between LLPS and chromatin compression has been demonstrated in many studies [33]. Moreover, phase-separated liquid condensates also control mRNA localization and processing [34–37]. The mechanisms of RNA distribution and processing in cells are important for the subsequent localization and function of proteins, highlighting the multistep nature of gene expression regulation [38]. LLPS is also involved in DNA damage response (DDR) through the formation of repair compartments by various DDR mediators [39–43]. Proper DDR is responsible for protecting the integrity and stability of the genome, and abnormalities associated with it can lead to oncogenesis [44,45]. It turns out that LLPS is associated with a wide range of diverse pathological processes, not only cancer-related, but also viral infections and neurodegenerative diseases [46–51]. Especially in the latter, LLPS is often a step preceding the pathological aggregation of disease-specific proteins [52].

Transcription factor 4 (TCF4) is a mammalian transcription factor (TF), also known as immunoglobulin transcription factor 2 (ITF2) or SL3-3 enhancer factor 2 (SEF2). TCF4 is a member of class I bHLH (basic helix-loop-helix) family [53]. These, also called E-proteins, are related to *Drosophila* Daughterless. In mammals, these proteins are TCF4, TCF3 and TCF12 [53,54]. The TCF4 C-terminal bHLH domain is responsible for dimerization and DNA binding [55,56]. The highly conserved basic (b) region composed of positively charged amino acids allows the protein to interact with the negatively charged Ephrussi box (E-box) DNA element (5’-CANNTG-3’) [55,57] in the promoters and enhancers of TCF4 responsive genes, such as the μE5 heavy and κE2 light chain immunoglobulin enhancers [58], an enhancer in the murine leukemia virus SL3-3 genome [59], the rat tyrosine hydroxylase enhancer [60], and the human somatostatin receptor-2 promoter [61]. In turn, HLH is involved in the formation of homo- and heterodimers with other bHLH TFs. After dimerization, the bHLH domains form a four-helix bundle with a hydrophobic core, where each bHLH monomer contacts an E-box half-site [55,62]. Depending on its bound partner in the heterodimer, TCF4 can act as a repressor [60,63,64] or activator [65–67] of transcription.

TCF4 exists in humans in many different isoforms due to alternative splicing - TCF4 mRNAs are expressed ubiquitously, but levels of different TCF4 isoforms are different in different tissues. Low-molecular-weight isoforms predominate in the brain: frontal cortex, cerebellum, and hippocampus, while those with higher molecular mass are predominant in the testis and lung [68]. All the isoforms have C-terminal bHLH domain, and they differ mainly in the length of N-termini [68] of TCF4, which contain regulatory domains. These domains allow TCF4 to regulate transcription independently of DNA [69–74]. In the longest TCF4 isoform (TCF4-B), these are two activation domains (AD1, AD2) and two repression domains, conserved element (CE), and repression domain (Rep). AD1 and AD2 are known to modulate transcriptional activity by interaction with transcriptional activator p300 [72,75] or repressors of transcription [76]. In turn, repression domain CE is located between AD1 and AD2 and can repress AD1 activity [77]. Rep falls between AD2 and bHLH and was shown to repress the activity of both AD1 and AD2 [78]. In addition to domains directly involved in transcriptional regulation, TCF4 contains various, partially overlapping localization signals - two nuclear localization signals (NLSs) and nuclear export signals (NESs) that control subcellular localization. NLS-1 with the putative activity of nucleolar localization signal (NoLS) overlaps with the CE domain at the N-terminal region and NLS-2 overlaps with NES-1 and NES-2 in the bHLH domain at the C-terminal region [68,79]. The first studies of human TCF4 indicated that it is localized in the cell nucleus [59], but later it turned out that only the longest isoforms are localized exclusively in the nucleus, while the shorter, lacking NLS-1 are present in both the nucleus and the cytoplasm [68]. Later studies have shown that, in most cases, TCF4 in neurons has a cytoplasmic localization [80], in line with the results of the tissue-specific distribution of individual isoforms [68], suggesting that the protein may have additional, non-genomic functionality. What is more, our recent studies have shown that, except for the bHLH domain, the remaining part of the shortest TCF4 isoform (TCF4-I ^-^) does not exhibit a stable fold [81]. This gives the N-terminal IDR of TCF4 the potential to gain a great range of conformations, enabling interactions of regulatory domains with a wide range of regulatory proteins. In contrast, a well-conserved basic part of bHLH interacts with a specific DNA sequence, anchoring TCF4 at the appropriate location in the genome. Together, these two important structural features of TCF4 have been proposed to allow it to act as a hub protein in central nervous system development [81,82].

TCF4 is essential for brain development, memory, and cognition. Mutations in *TCF4* gene have been linked to schizophrenia, autism-spectrum intellectual disability, and Pitt-Hopkins syndrome (PTHS) [83]. PTHS is a rare genetic disorder characterized by varying clinical severity, including intellectual disability, motor impairments, and autistic features [84,85]. It is caused by the wide range of mutations in *TCF4*, which may lead to its haploinsufficiency or mutations of TCF4 protein [86,87]. These are usually point mutations located in TCF4 bHLH [63,64,86–89] and most of them were observed to decrease DNA binding ability and TCF4-mediated transcription activation [64,89,90]. Interestingly, some of these mutants exhibit different subcellular localization patterns than the wild-type TCF4, which is distributed equally. These mutants were observed to form nuclear puncta, but the mechanism of assembly and material properties of these puncta were not determined [64,89,90]. Notably, there have been reports of associations between TCF4 mutations occurring outside the bHLH domain and schizophrenia. Common variants in the human *TCF4* were among the first genes to reach significance in genome-wide association studies of schizophrenia [91], and rare coding *TCF4* variants outside of the bHLH domain were identified in individual patients with schizophrenia by deep sequencing [92,93].

In this study, we present the results of LLPS analyses of TCF4 isoform I^-^ and the intriguing influence of DNA on the formation of TCF4 condensates. TCF4-I^-^ is the shortest human TCF4 isoform, predominant in the brain [68]; what is more, the only one with determined molecular properties [81]. We demonstrate that purified TCF4 undergoes LLPS, and the DNA modulates this process. We also determined that TCF4 condensates can undergo maturation, and that the dynamics of TCF4 within these condensates change. Based on these findings, we propose that LLPS of TCF4 is its natural property that enables TCF4-mediated transcription burst, but under some conditions, TCF4 LLPS may be disease-related. We suggest that the altered cellular localization pattern of some PTHS-associated TCF4 mutants may be related to affected LLPS, which leads to the shutdown of TCF4 function through the formation of pathological, solid-like condensates. We believe that our study will partially fill the knowledge gap, since phase separation has recently been proposed as a mechanism of pathogenicity for disease-linked mutations [94].

## 4. Materials and methods

### 4.1 In silico analysis of TCF4

In this article, all experiments, with the exception of cell imaging, were conducted using the human TCF4 isoform I^-^ (UniProtKB P15884-16), hereafter referred to as TCF4 in subsequent sections.

The analysis of the TCF4 sequence employed bioinformatics tools with default settings. Analyses of the TCF4 propensity to undergo LLPS were carried out using independent predictors, FuzDrop [95– 97], PScore [98], ParSe [99] and catGRANULE [100].

### 4.2 Preparation of TCF4 for LLPS studies

TCF4 isoform I^□^ was expressed, purified and labeled as described before [81]. For localization studies in COS-7 cells, deletion mutants of the TCF4-B^+^ isoform (UniProtKB Q60722): CFP-TCF-B^+^ /1-550 and CFP-TCF-B^+^ /1-601 cDNA vectors were prepared analogically to the preparation of the YFP-TCF-B^+^ /1-550 and YFP-TCF-B^+^ /1-601 cDNA vectors described previously [79]. Also, COS-7 cells culture condition and transfection was performed as described previously [79].

### 4.3 Laser scanning confocal microscopy imaging

As described previously [79], to prepare cells for confocal microscopy imaging, before cells passage, 0.17-mm-thick round glass coverslips (Menzel) were added to 2-cm diameter Petri dishes and submerged in DMEM. Next, 24h after transfection, the coverslips with cells were transferred onto a steel holder and mounted on a microscope stage. The medium was replaced with 1⍰mL of DMEM/F12 without phenol red, buffered with 15⍰mM HEPES (Sigma) and supplemented with 2% FBS (Sigma). The cell culture temperature was stabilized at 37⍰°C by using a microincubator (Life Imaging Services Box & Cube). Images of the CFP labelled proteins were acquired using a Leica TCS SP5 II confocal system equipped with argon and helium neon lasers and a 63 × oil objective lens (NA: 1.4). CFP was excited using 458⍰nm light and the emitted fluorescence was observed at a range of 470-600⍰nm. To test the character of the liquid condensates, 10% 1,6-hexanediol was added to the plates with cell culture.

To initiate LLPS *in vitro*, TCF4 (Alexa Fluor 488-labeled TCF4 with unlabeled TCF4 in a 1:20 ratio) in buffer S was mixed by pipetting with appropriate amounts of NaCl, to achieve a final protein concentration of 1 mg/mL. To investigate the effect of NaCl on TCF4 LLPS, the liquid phases were visualized 3 and 5 min after incubation without a cover. For condensate maturation studies at 500 mM and 750 mM NaCl, condensates were imaged at intervals for up to 5 h in a solution placed in Press-to-Seal silicone isolator (9 mm diameter, 0.5 mm deep, Invitrogen), covered with a glass slide. Coalescence was observed at 500 mM NaCl after 3 min of incubation with a cover (as described above) and recorded at 5 s intervals. Measurements were made in 10 µL or 4 µL condensates at 23 °C on unpassivated microscopic slides. Images were acquired using an inverted laser scanning confocal microscope (Axio Observer LSM 780, Zeiss), with a MBS 488 beam splitter and a BP 495-555 nm filter, with the excitation at 488 nm using an argon ion laser and a Plan-Apochromat 63 × (NA 1.40) oil DIC M27 objective.

### 4.4 Fluorescence recovery after photobleaching

Fluorescence Recovery After Photobleaching (FRAP) experiments in cells were performed using FRAP Wizzard of Leica Application Suite Advanced Fluorescence (LAS AF). The selected point ROI located in the puncta of the expressed in COS-7 cells nucleus CFP-TCF4-B^+^ /1-601 mutant protein, was bleached by 2-10 seconds with 100% laser power at 458 nm. Fluorescence recovery was monitored for at least 4 minutes.

*In vitro* FRAP on purified TCF4 was performed on two to five droplets per each condition that were bleached with the 100% laser power. Fluorescence recovery was monitored for at least 60 s. Data analysis was performed on both individual and averaged runs using FRAPAnalyzer 2.1.0, accounting for the photobleaching (reference) and background signal. The data were normalized, and a one exponential recovery model was fitted:

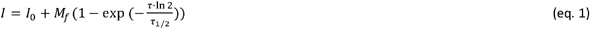

where: *I*, fluorescence intensity; *I*_*0*_, residual fluorescence after bleaching; *M*_*f*_, mobile fraction; *τ*, time after bleaching; *τ*_*1/2*_, recovery half-time. The *τ*_*1/2*_ and *M*_*f*_ values were determined as the arithmetic mean from individual runs, with the standard deviation as the experimental error.

The kinetic rate constant, *k*, for FRAP was calculated as:

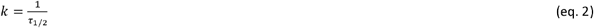

The Akaike Information Criterion (AIC) was used to select the appropriate numerical model to analyze the dependence of the FRAP results, *M*_*f*_ and *τ*_*1/2*_, on the experimental conditions, *i.e*., NaCl concentration, *c*, and incubation time, *t*. In the case of mobile fraction, a linear function with the slope fixed to 0 was chosen after comparing it with a linear model in which both parameters were free. This function was then fitted to *M*_*f*_(*t*) (AIC = 0.85) and to *M*_*f*_(*c*), the latter measured after 3 min (AIC = 0.99) and 5 min (AIC = 0.99) of incubation.

A two-parameter exponential function:

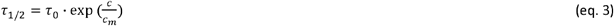

was chosen after the comparison with a linear function, and fitted to the *τ*_*1/2*_ *vs. c* data obtained after *t* = 3 min (AIC = 1) and 5 min (AIC = 0.99) of incubation. Here, *τ*_*0*_ is *τ*_*1/2*_ at NaCl concentration *c* = 0, and *c*_*m*_ is the *c* value at which the *τ*_*1/2*_ increases by a factor of *e*.

In the case of the *τ*_*1/2*_(*t*) data, a one-phase growth function proved to be the minimal model with a physical interpretation that could explain the condensate maturation process, as follows:

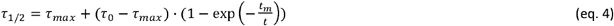

where *τ*_*0*_ is *τ*_*1/2*_ at *t* = 0; *τ*_*max*_ is the plateau value of *τ*_*1/2*_; *t*_*m*_ is a characteristic time constant of condensate maturation.

All fits were performed with the data points weighted by the experimental errors (*w*= *sD*^-2^). Error calculus was performed according to propagation rules based on the partial derivatives of the variables.

### 4.5. Widefield microscopy imaging

For phase diagrams preparation, TCF4 samples in buffer S were concentrated to 3 mg/mL using the Amicon Ultra-4 Centrifugal Filter Unit (Merck/Millipore; molecular weight cut-off 30.0 kDa). The concentrated solution was used to prepare a series of working solutions. The TCF4 samples were analyzed at various concentrations ranging from 0.25 to 1.5 mg/mL and NaCl concentration 100 - 1000 mM without additives and with 20 μM dsE-box. For investigations of the influence of NaCl concentration on TCF4 aggregation and differentiation of aggregates and liquid condensates, TCF4 labelled with Alexa Fluor 488 was used. The samples in buffer S were prepared to obtain final concentrations of 1.5 mg/mL TCF4 (labeled:unlabeled TCF4 1:20 ratio) and 800 mM to 1.5 M NaCl.

For condensates dissolving experiments, the samples in buffer S were prepared to obtain final concentrations of 1 mg/mL TCF4, 700 mM NaCl and 20 μM DNA (dsE-box, ssE-box, double stranded non E-box), or 0.5 mg/mL Alexa Fluor 488-labeled TCF4 (labeled:unlabeled TCF4 1:20 ratio), 150 mM NaCl, 5% PEG 8000, and 10 μM FAM-dsE-box DNA. The DNA was previously dissolved in buffer S and with pH adjusted to 7.5. The sequences of DNA used were as follows:

E-box: CCGGTCACGTGTCCTA, where the E-box canonical sequence is in italics

non E-box: AGTGCTGAGCCTGAAC

Widefield microscopy was performed using an Axio Observer 7 (Carl Zeiss) inverted microscope with 100 × oil and 1.3 numeric aperture objective lenses. The samples were incubated for 5 min, then 10 μL volume analyzed without cover at RT. Differential interference contrast (DIC) and fluorescence images were collected with an Axiocam 305 color camera (Carl Zeiss).

### 4.6. Turbidity measurements

Turbidity measurements of TCF4 without additives for phase diagrams were performed on the same set of samples as for widefield microscopy imaging, in 2 μL of the sample. The absorption spectra in the range of 220-700 nm were recorded using NanoDrop 2000c. Then the differences of the spectra, especially in A_280_, A_340_ and A_600_ of samples at different NaCl concentrations were determined.

### 4.7. Fluorescence anisotropy measurements

All the samples were prepared to a final volume of 60 μL and contained constant concentration of 0.5 mg/mL TCF4 (Alexa Fluor 488-labeled TCF4 with unlabeled TCF4 in a 1:20 ratio) and different concentrations of PEG 8000 (0 - 10% w/v) in buffer S. Fluorescence anisotropy measurements were performed on an FP-8300 spectrofluorometer (Jasco) equipped with excitation and emission polarizers and with a Peltier temperature controller (EHC-813). QS High Precision Cell cuvettes (105– 251-15–40; Hellma Analytics) were used. All the samples were monitored after 5 min of PEG addition, at an excitation wavelength of 493 nm and emission wavelength of 516 nm, with 5-nm bandwidths. The changes in fluorescence anisotropy were monitored at 25 °C. Anisotropy (*r*), including the instrument G factor (1.1219), was calculated using the equation:

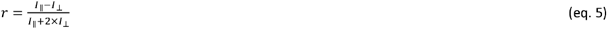

where *I*_‖_ and *I*_⊥_ are the components of the fluorescence intensity that are parallel and perpendicular to the electric vector of the excitation light, respectively [101].

The same set of samples was used to further confirm the condensates’ existence using widefield fluorescence microscopy.

## 5. Results

### 5.1. TCF4 deletion mutant lacking C-terminal region of bHLH forms condensates in the cytoplasm

In our previous work, we performed detailed studies on the localization of different TCF4 deletion mutants in COS-7 and N2a cells [79]. In contrast to the CFP-tagged mutant lacking the entire bHLH domain (CFP-TCF4-B^+^/1-550), which was distributed almost evenly throughout the nucleus, except for the nuclei (Fig. 1A), we observed that the mutant lacking the last 69 amino acid residues of TCF4, mostly encompassing the C-terminal region of the bHLH domain (CFP-TCF4-B^+^ /1-601), exhibited a specific punctate pattern within the nucleus (Fig. 1A). Interestingly, this pattern was similar to previously observed patterns of TCF4 bHLH mutants linked to PTHS [64,79,90]. The pattern also resembled that presented by proteins capable of phase separation, whose ability to separate and interact has been linked to the development of neurodegenerative diseases [49]. To verify hypotheses about the liquid nature of the observed puncta in nuclei we also performed the FRAP experiment. The fluorescence recovery was so quick that we could observe a small darker area only immediately after ROI bleaching, even when using longer bleaching times (up to 10 s) than presented on Fig. 1B, C. In contrast, we observed an immediate increase in the fluorescence area of the punctus including the selected ROI (Fig. 1C). These results have led us to study the ability of TCF4 to be able to form condensates through the LLPS more extensively. We decided to test the effect of 1,6-hexanediol, known to dissolve condensates formed due to hydrophobic interactions [102], on the expressed CFP-TCF4-B /1-601 TCF4 mutant. Surprisingly, the addition of this compound resulted in the rapid, total disappearance of the puncta in a very short time, less than 2 seconds (Fig. 1D). Since the protein’s propensity to LLPS is affected by many factors, all of which cannot be controlled, especially in a living cell, we decided to first carry out studies on the purified TCF4 protein, which is the subject of the work presented here.

**Figure 1.**
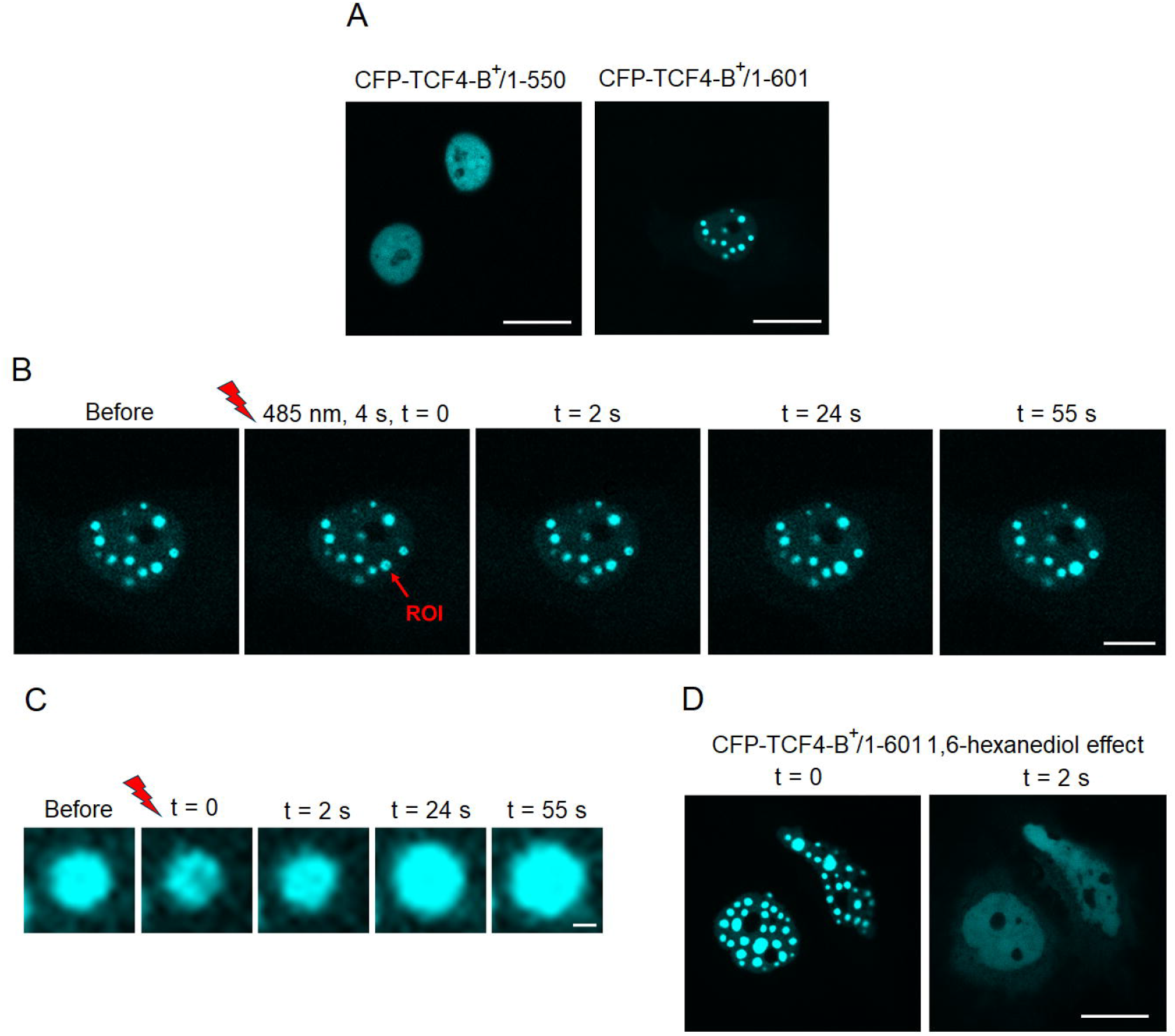

### 5.2 *In silico* analyses indicate that N-terminal part of TCF4 may be responsible for LLPS

In our studies, we focused on the shortest isoform of TCF4, I ^-^, which is predominant in neurons [68]. This is the only form of TCF4 with experimentally determined molecular properties so far. Our previous studies have shown that except for the bHLH domain, the protein is completely disordered [81]. Considering that IDRs play a huge role in the LLPS process [4–9] we decided to test TCF4 for its propensity to undergo LLPS. We began with *in silico* analyses choosing a set of various LLPS predictors: FuzDrop [95–97], PScore [98], ParSe [99], and catGRANULE [100].

The analyses performed using FuzDrop and PScore were in general agreement showing that the C-terminal fragment of TCF4, which mainly contains bHLH, is not likely to undergo LLPS, while the long, disordered N-terminal TCF4 fragment [81] has a high probability of undergoing LLPS (Fig. 2A, B). The main difference was that the PScore analysis indicated a disordered fragment covering amino acids from 325 to 350 as having no LLPS propensity (Fig. 2B), which is inconsistent with the FuzDrop analysis result (Fig. 2A). Overall LLPS propensity scores classified TCF4 as very likely to undergo LLPS (overall PScore: 2.65 and FuzDrop pLLPS: 0.9998, respectively). In contrast, the results of the ParSe analysis suggested a more mosaic nature of TCF4 - alternating disordered regions with LLPS propensity, without LLPS propensity, and those that can fold into a stable structure (Fig. 2C). The longest region with LLPS potential was determined as the residues 1-169 (and the shorter region, residues 194-227), while the regions predicted as able to form stable structures were residues 170- 188 and 228-247, which do not correspond to any known TCF4 partner-interacting regions, and the region of 354-408, which completely covers bHLH domain. The results obtained with catGRANULE also suggested a high propensity of TCF4 to undergo LLPS (an overall propensity score of 1.287), but in contrast to other predictors, regions with the higher LLPS potential included residues 154-218 and 258-307 (Fig. 2D). These analyses not only suggest the probable role of TCF4 in LLPS, but also highlight the importance of the concurrent use of different predictors that operate under different principles.

*In silico* analyses suggested that TCF4 is highly likely to undergo LLPS. The regions identified as the most probable to be involved in LLPS, were regions previously classified as IDRs [81]. As even advanced *in silico* analysis cannot provide evidence of LLPS ability, experimental studies were necessary to verify TCF4 ability to induce spontaneous LLPS.

**Figure 2.**
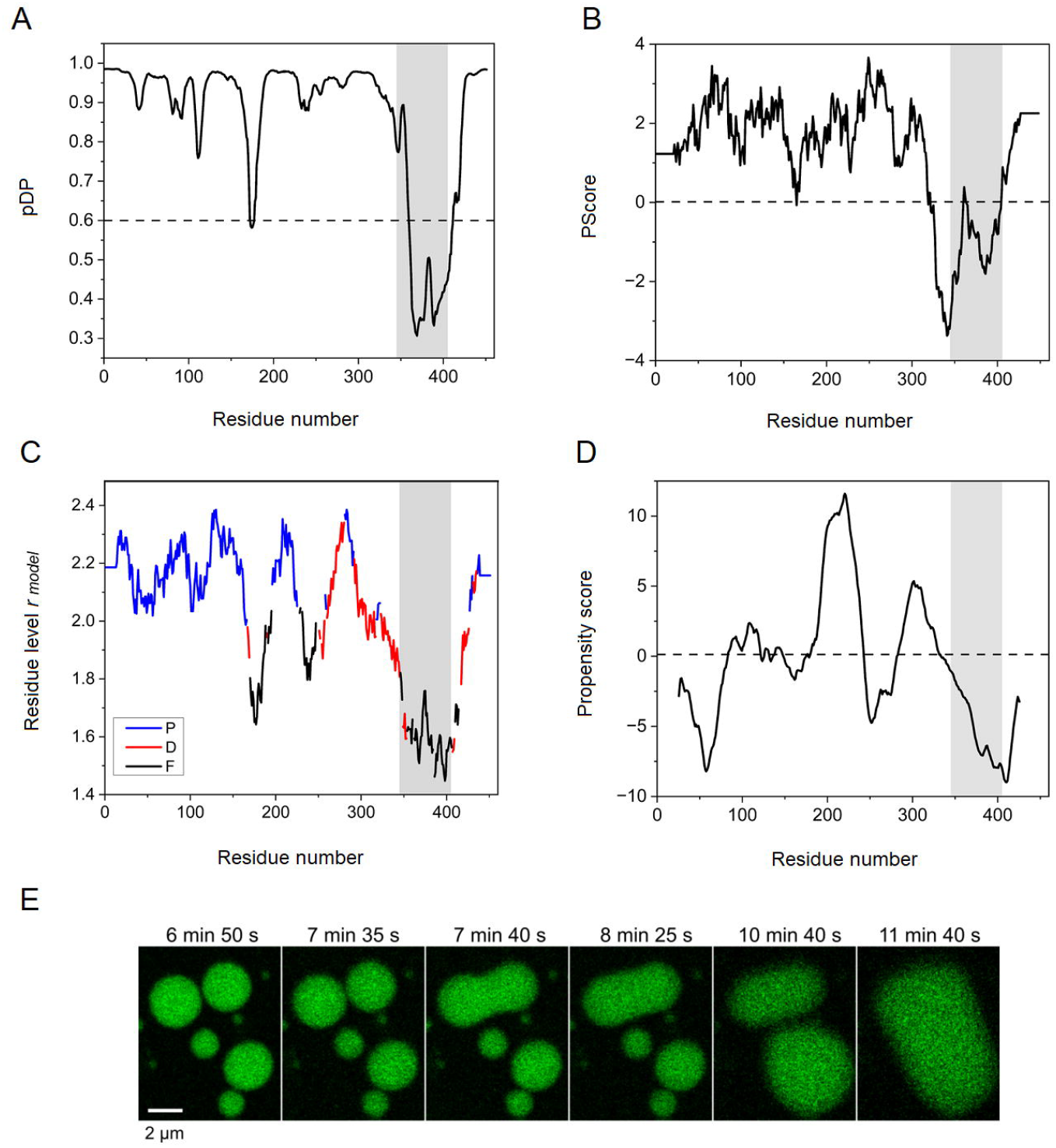

### 5.3. Increase of ionic strength leads to TCF4 phase separation *in vitro*

While studying the effect of environmental conditions on TCF4, we discovered that increasing the ionic strength of a sample by raising the concentration of NaCl results in phase separation of TCF4 (including ∼ ∼5% of fluorescently labeled protein). This effect was evident at the NaCl concentration range of 400 to 900 mM (Fig. 3A). The dynamic fluorescent assemblies originating from TCF4 were observed as condensates that were mobile and had the ability to coalesce (Fig. 2E, Supplemental Video), suggesting their liquid-like nature.

**Figure 3.**
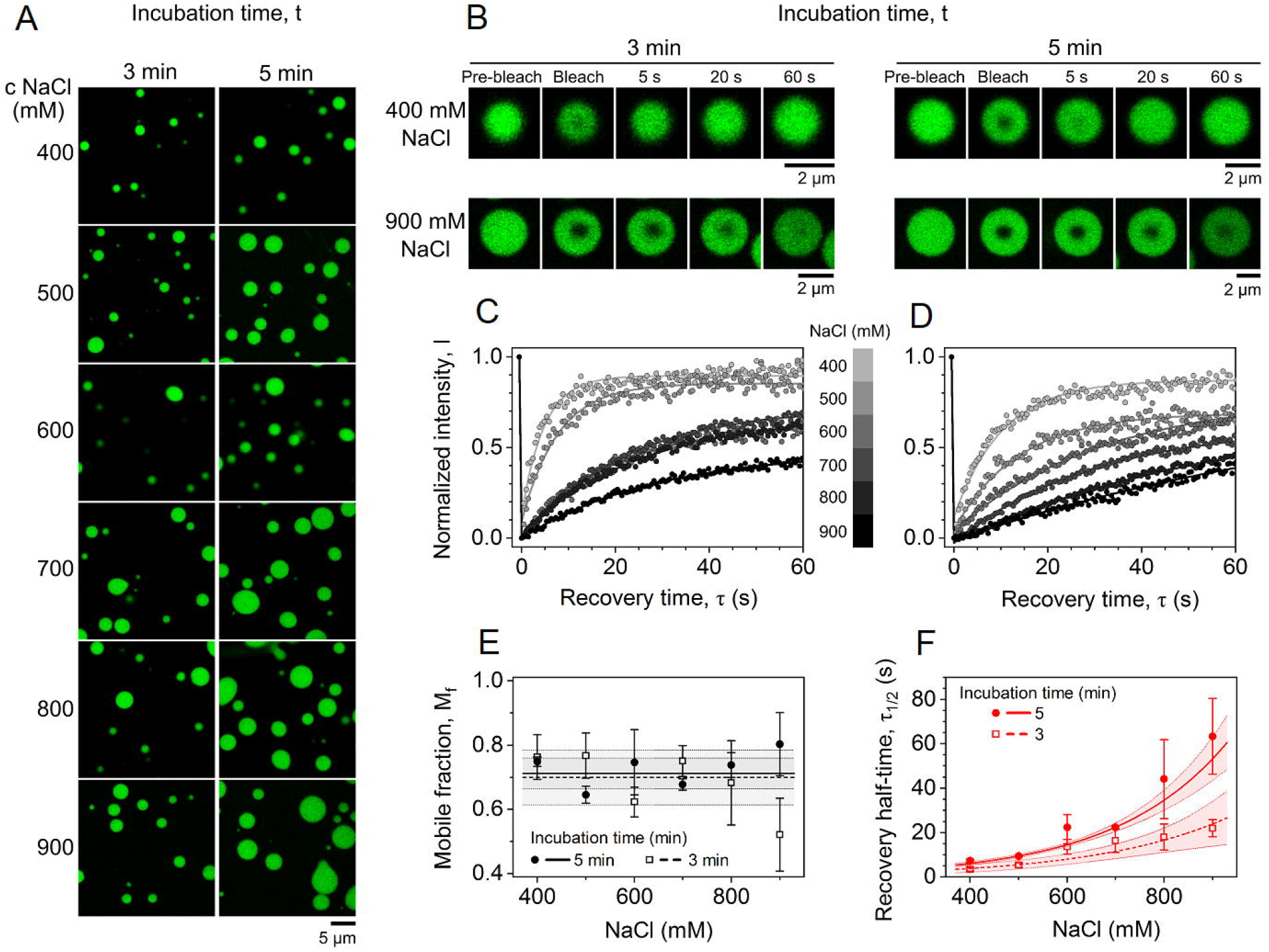

Therefore, we aimed to characterize the liquidity of those TCF4 assemblies across the range of increasing NaCl concentrations corresponding to the phase separation window. To this end, FRAP experiments were performed on condensates formed at 1 mg/mL TCF4 after 3 and 5 minutes of incubation with the salt (Fig. 3B-D). Regardless of the concentration of NaCl and the incubation time, the condensates retained their liquid-like properties, as evidenced by a high and constant mobile fraction of approximately 70% across all conditions (Fig. 3E). However, an exponential increase in the recovery half-time, τ1/2, was observed with increasing NaCl concentration (Fig. 3F). Interestingly, the characteristic concentration values, *c*_*m*_, associated with this process, determined from eq. 3 (see Materials and methods section), were statistically comparable at both incubation times, with *c*_*m*_ = 270 ± 40 and 231 ± 11 mM, for 3 and 5 minutes of incubation, respectively.

The reduction in the diffusion rate constant of TCF4 (Table S1) within condensates at higher ionic strengths likely stems from salt-mediated charge screening, which diminishes electrostatic repulsion and promotes stronger short-range hydrophobic interactions and multivalent cross-links. Consequently, the condensate’s internal network becomes denser, which raises its viscosity and hinders molecular diffusion. Notably, the recovery half-times measured after 5 minutes of incubation were consistently longer than those measured after 3 minutes at equivalent NaCl concentrations (Fig. 3F). This suggests that, over time, the properties of TCF4 condensates may change over time, leading to their hardening through a process known as condensate maturation [103]. This observation prompted us to further investigate the impact of incubation time on TCF4 condensate material properties.

### 5.4. Investigation of the TCF4 condensates aging process

To study the effect of time on TCF4 condensates, these formed at 11mg/mL TCF4 in 500 mM NaCl were observed over a 5-hour period. With increasing incubation time, the area occupied by TCF4 condensates gradually increased, likely due to progressive condensate coalescence and sedimentation (Fig. 4A). The FRAP analysis (Fig. 4B, C) revealed that the recovery half-time *τ*_*1/2*_ increases asymptotically with incubation time *t* (Fig. 4D), with a characteristic time *t*_*m*_ of 32 ± 8 min and maximal *τ*_*1/2*_ (*τ*_*max*_) of 20.0 ± 0.8 s, according to the eq. 4 (see Materials and methods section). Surprisingly, the mobile fraction remained unchanged despite increasing incubation time. This indicates a maturation process in which the condensates become more viscous but do not transition into a solid or gel-like state that immobilizes a fraction of the molecules. Such behavior implies that the internal network of multivalent interactions and hydrophobic contacts strengthens over time, restricting diffusion rates (Table S2) without fully arresting molecular mobility. Consequently, TCF4 condensates maintain their liquid-like properties within the examined time range, lasting up to 5 hours. Analogous analyses were carried out for TCF4 condensates at 750 mM NaCl, in which similar trends were observed (Fig. S1, Table S2).

**Figure 4.**
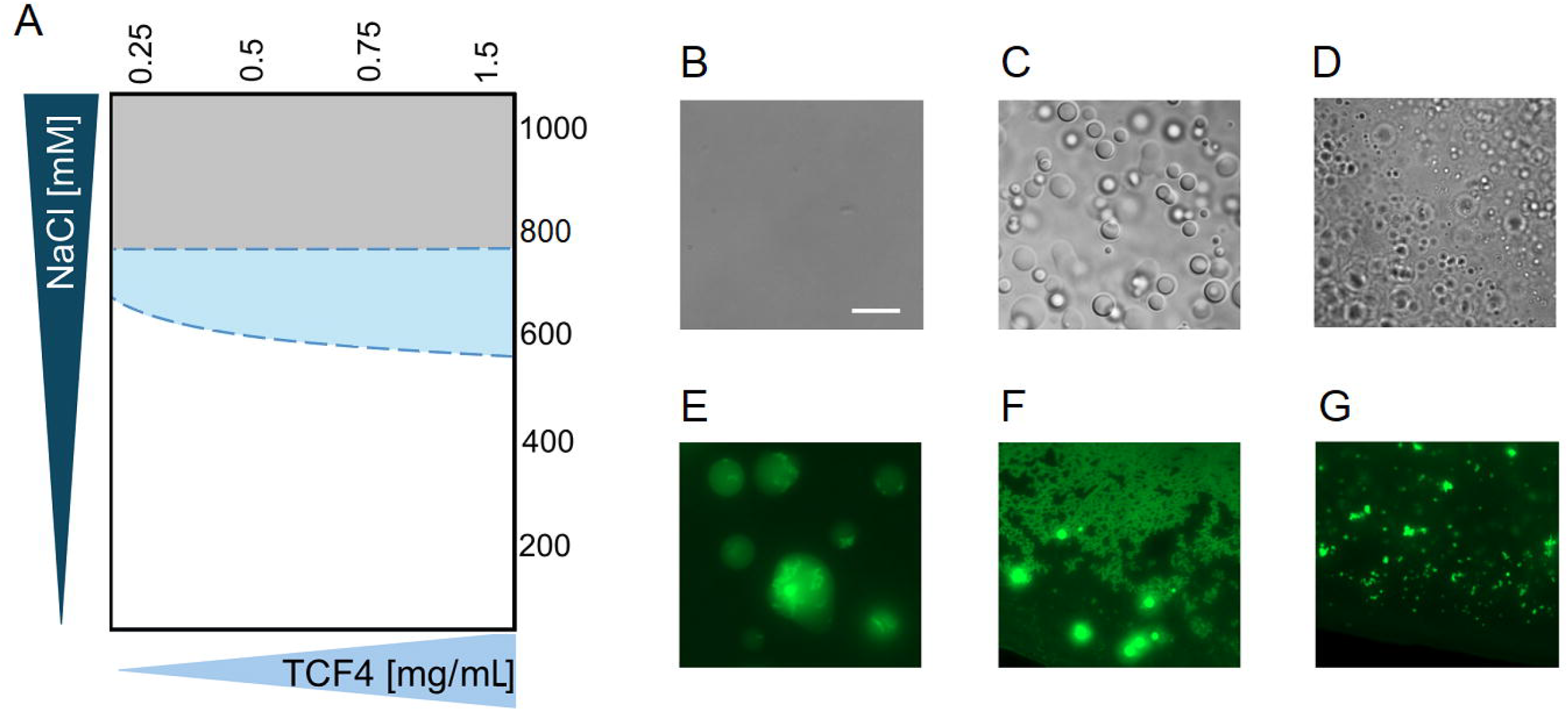

### 5.5. High NaCl concentrations lead to TCF4 aggregation

We performed screening of different NaCl concentration effects on unlabeled TCF4 at different concentrations. Analyses were performed using widefield microscopy imaging and turbidity measurements. We analyzed TCF4 at concentrations ranging from 0.25 to 1.5 mg/mL in a buffer containing 100 – 1000 mM NaCl. Using microscopic imaging, we observed homogenous solutions for TCF4 samples containing NaCl from 100 mM to 500 – 600 mM (depending on TCF4 concentration) (Fig. 5A; white area, Fig. 5B, Table S3). Samples containing at least 600 – 700 mM NaCl, depending on protein concentration, revealed the presence of TCF4 spherical condensates (Fig. 5A; blue area, Fig. 5C Table S3). Analysis of TCF4 samples in higher salt concentrations (at least 800 mM NaCl) revealed that there are different types of TCF4 assemblies present in the sample (Fig. 5A; gray area, Fig. 5D Table S3). Some of them seemed to be droplet-like in shape and mobile, while others were static with an irregular shape. The recorded absorption spectra of the analyzed samples were generally consistent with microscopic observations. Samples that were homogeneous in microscopic observation had typical absorption spectra that were reproducible at different salt concentrations. Samples for which microscopic images showed the presence of typical droplets were characterized by altered absorption spectra. These alterations were observed as changes in A_280_, generally with a simultaneous increase in A_340_ and A_600_, indicative of sample turbidity characteristic for LLPS. Interestingly, samples that contained different types of TCF4 assemblies revealed ordinary absorption spectra, the same as samples containing a homogeneous solution of TCF4 (Fig. S2).

**Figure 5.**
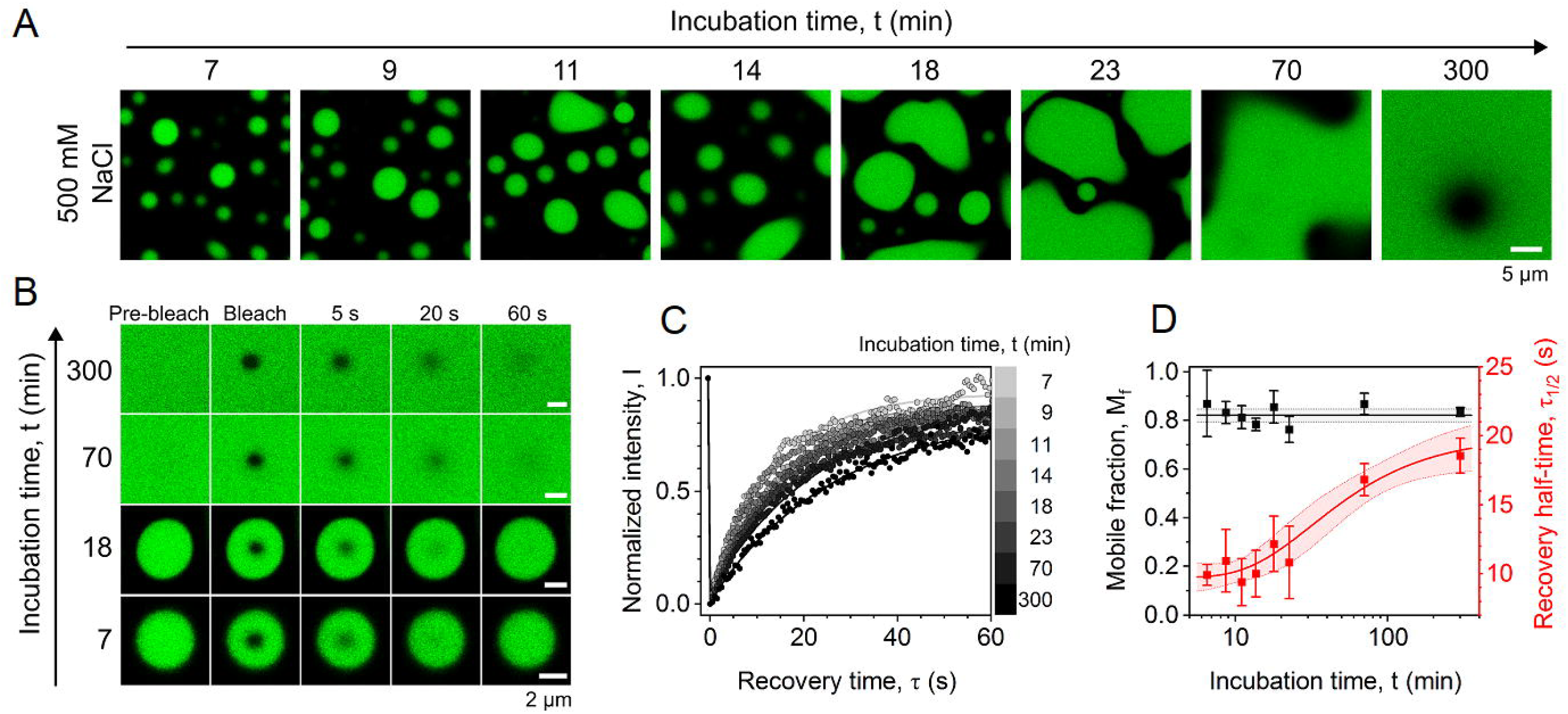

These data provided information on the NaCl concentrations at which TCF4 undergoes phase separation slightly different from those obtained for the labeled protein. This discrepancy may be due to experimental conditions or fluorescent dye, highlighting the well-known conviction that fluorescent dyes are not inert in experimental results. However, since DIC did not enable us to accurately distinguish droplets from aggregates, we decided to once again use fluorescence microscopy. We analyzed TCF4, including ∼5% of fluorescently labeled protein, at 1.5 mg/mL at different, high NaCl concentrations. It revealed coexisting droplets and aggregates of TCF4. Irregular shape aggregates inside the TCF4 droplets were observed at NaCl concentration of 800 mM (Fig. 5E), while independent aggregates and droplets were present in the sample containing 1 M NaCl (Fig. 5F). 1.5 M NaCl led only to the formation of TCF4 aggregates and droplets were not observed (Fig. 5G).

### 5.6. DNA inhibits NaCl-dependent LLPS of TCF4

Since TCF4 is a transcription factor that binds to E-boxes in the promoters and enhancers of responsive genes [54,58–61,104], we tested the effect of DNA on TCF4-mediated LLPS induced by increased ionic strength. For this purpose, we prepared samples containing TCF4 at 0.25-1.5 mg/mL, different NaCl concentrations (100-1000 mM), and 20 μM dsE-box. We did not observe TCF4 condensates under the conditions we studied, which suggests that DNA prevents TCF4 from LLPS. Next, we examined the effect of DNA on existing TCF4 condensates. For this purpose, we initiated TCF4-driven LLPS by increasing the NaCl concentration, and then added dsE-box to the preformed TCF4 condensates (Fig. 6A). The solution became homogeneous, indicating that dsE-box not only prevents TCF4 from NaCl-dependent LLPS, but also leads to the dissolving of existing TCF4 condensates. To determine if the dissolving effect of DNA on TCF4 LLPS is due to specific sequence binding or rather the presence of DNA in general, we performed analogous analyses using ssE-box and non-specific dsDNA (Fig. 6B). The effect on TCF4 LLPS was the same as in the case of dsE-box: the TCF4 condensates were not observed after the addition of DNA. A control experiment using buffer S (Fig. 6B) revealed that the dissolution of TCF4 condensates was not due to sample dilution.

**Figure 6.**
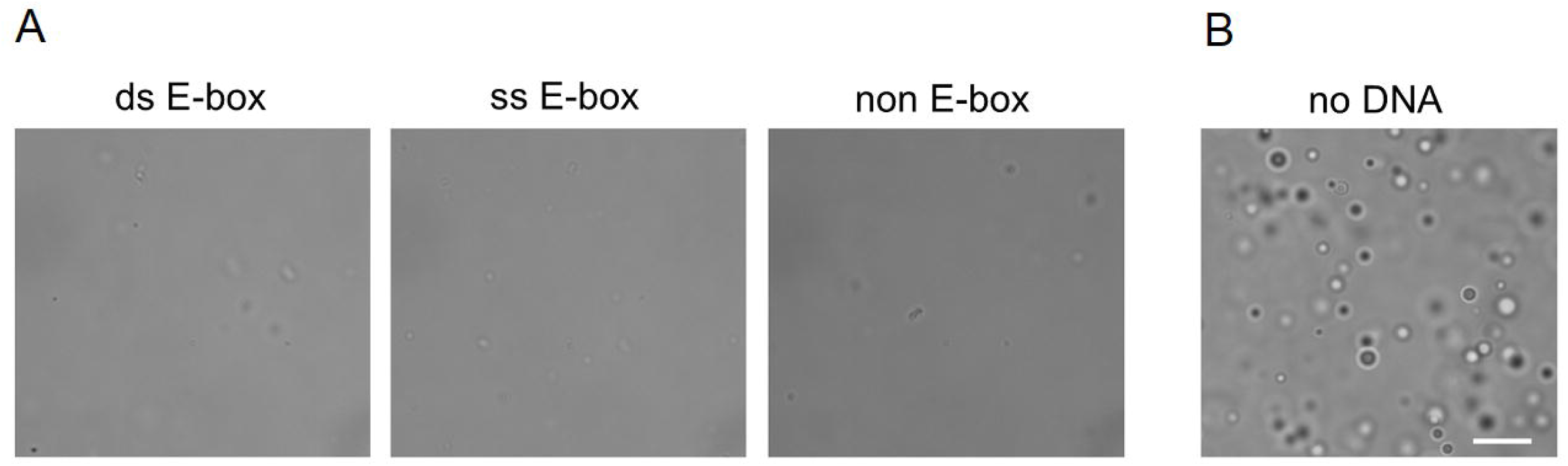

### 5.7. TCF4 forms condensates under molecular crowding conditions

We also determined the effect of molecular crowding on TCF4. Homogenous solution of TCF4 (including ∼5% of fluorescently labeled protein) at 0.5 mg/mL in buffer S was analyzed in the presence of varying PEG 8000 concentrations (Fig. 7). The anisotropy measurements revealed that its value increases with the increasing concentration of PEG, indicating formation of fluorescent assemblies with their size directly correlated with PEG concentration (Fig. 7F). This was confirmed by microscopy imaging, where we have observed the formation of labeled TCF4 assemblies for the concentrations of PEG in the range of 3-10% (Fig. 7A-E). Then we tested the effect of E-box on TCF4 condensates formed as a result of molecular crowders addition (Fig 7G, H). For this purpose, we used unlabeled TCF4 and FAM-labeled dsE-box. We observed that, at the conditions studied, the DNA not only has no dissolving effect on TCF4 condensates, but it also diffuses into them. This result was contrary to the dissolving effect of DNA on NaCl-dependent TCF4 condensates.

**Figure 7.**
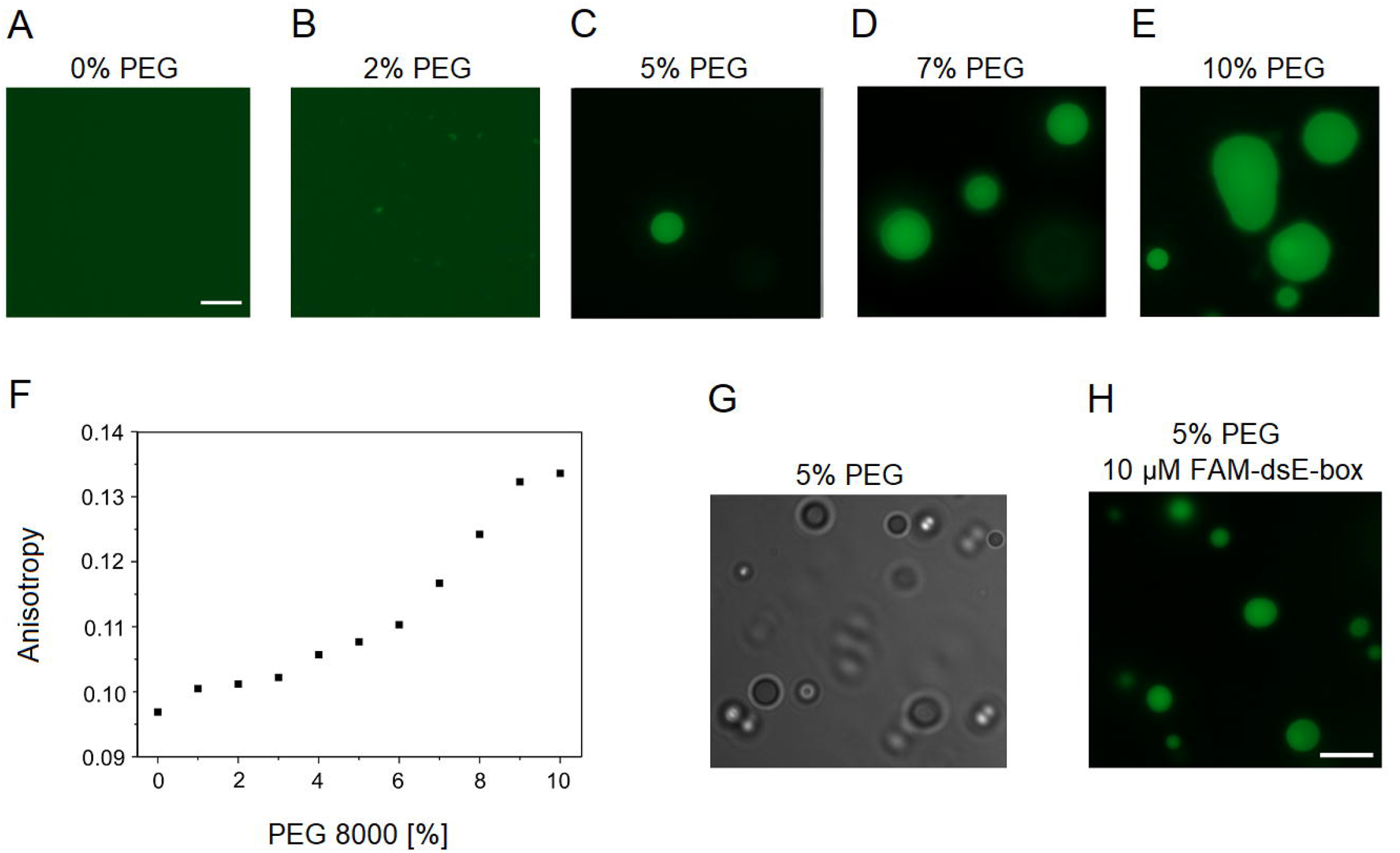

## 6. Discussion

In this study, we show that the TCF4 isoform I^□^ undergoes phase transition and present the results of analyses of the intriguing influence of DNA on the formation of the TCF4 condensates. The isoform I ^□^ is deficient in amino acid residues 1-216 in comparison with the canonical isoform B^□^ (UniProt ID P15884-1), and consequently, it lacks the region comprising AD1, the N-terminal NLS, and the CE repressor domain, but it contains the entire bHLH domain [68] (Fig. S3). We recently showed that approximately 90% of TCF4-I ^□^ consists of disordered regions [81]. Only the HLH motif responsible for dimerization possesses a well-ordered fold. In addition, the basic fragment (b), which is important for E-box binding, folds upon binding. These dual structural properties may explain the ability of involvement of TCF4 to act as a “hub” for the regulation of transcription [81,82]. In this context, we previously proposed that TCF4 functions in two independent modes. The first is related to the function of the bHLH domain and depends on the recognition and binding of the protein to specific DNA sequences, resulting in the TCF4 being anchored to the appropriate genomic location. In the other mode, TCF4 IDRs act as a flexible platform for multiple interactions with other transcription-related proteins, independently of DNA binding [81].

The intrinsically disordered character is particularly preferred for proteins with regulatory activity at the crossroads of signaling pathways, as high flexibility allows them to interact with multiple partners in a context-dependent manner [105]. Those interactions often involve many weak, transient self-interactions or interactions with nucleic acids. This may lead to the formation of multicomponent assemblies, known as MLOs [1,10,11], in the LLPS process [4–9]. Importantly, TCF4 exists as many isoforms, generated via alternative splicing, with tissue-specific expression pattern [68], and these isoforms differ in the length of N-terminal IDR. Interestingly, alternative splicing that results in the existence of isoforms with different IDR lengths has been proposed as a mechanism for regulating LLPS of the RNA-processing prion-like protein hnRNPDL [106].

Regulatory proteins, including TFs, are highly dynamic and interact transiently with DNA [107,108], searching through the genome while non-specifically bound to DNA until they find specific DNA sequences to which they bind with high affinity [109–111]. Thus, the interaction of TFs with DNA includes short-lived non-specific interactions, and the long-lived specific interactions [112–114]. While specific binding DNA requires direct hydrogen bonds and van der Waals forces between the TF and DNA base pairs [115], non-specific DNA binding relies on electrostatic interactions of the basic residues of a TF with the negatively charged DNA backbone [111]. These two modes of DNA binding engage two distinct TFs structural properties – DNA binding domain binds to specific DNA sequence, preceded by non-specific binding and search through the genome, which engage TFs IDRs. This statement is supported by deletion and mutation studies of DNA binding domains of TFs, which resulted in a loss of the specific, but not non-specific binding ability [112,116–118]. Thus, TF IDRs may not only lead to its self-interactions which promotes phase separation [1,119], but also non-specific DNA binding.

Based on an *in silico* analysis, TCF4 has been previously suggested to undergo LLPS, but this has not been confirmed to date [94]. Importantly, most of amino acid residues that are linked to the PTHS-associated point mutations of human TCF4 isoform B ^+^[63,89,120–122] interact directly with the DNA (PDB OD03, OD04, [104]). These are as follows: R569, R576, R578, R580, N585 and K607. In our study, TCF4-B^+^ /1-601 presented unique punctate pattern in the nucleus of analysed COS-7 cells (see Fig. 1) similarly to Forrest et al. [64] experiments in COS-7 and SH-SY5Y neuroblastoma cells, where the mutants R578P, R580W, and to a lesser extent A614V formed a small, spherical puncta, whose localization differed markedly from the distribution of the wild-type TCF4. Similarly, altered distribution of TCF4 mutants R569W and N585D were observed by Sirp et al [90]. According to our observations, TCF4-B^+^ /1-601 variant lacking the C-terminal fragment containing K607 and A614 was also present in the puncta in the nucleus. That might be connected to the impaired DNA binding due to missing DNA-binding residue (K607) or conformational changes caused by the loss of protein C-terminus. Using FRAP experiments and testing 1,6-hexanediol impact on the puncta in the nucleus resembling the PTHS-linked pattern of our mutant, we documented the liquid nature of formed spherical condensates. These findings support the hypothesis that TCF4’s capacity to form condensates is associated with its intrinsic propensity for LLPS, a process that requires strict regulatory control, as its dysregulation—particularly in mutant variants—may contribute to the pathogenesis of PTHS. For this reason, we believe that characterization of TCF4 ability to undergo LLPS and factors influencing this process is an important step toward understanding the nature of PTHS development.

Our *in vitro* analyses on purified protein revealed that TCF4 undergoes LLPS, forming droplets in a NaCl concentration-dependent manner. Droplets fuse, and TCF4 molecules have high mobility within condensates. Importantly, LLPS driven by TCF4 was not only inhibited, but also reversed by DNA addition. Our analysis revealed that it is not linked to the specific DNA sequence binding, but rather DNA presence in general. We propose that TCF4 accumulates in large amounts in condensates, which ensures that at the proper conditions, e.g. after binding a transcription activator, enough TCF4 molecules can dissociate from the condensate and be involved in a rapid and efficient transcription process. We hypothesize that in the absence of DNA TCF4 uses its IDR as LLPS platform (Fig. 8). In presence of DNA, TCF4 IDR preferentially interacts with DNA to search for specific sequence, which leads to dissolve of TCF4 condensates. It is worth highlighting that in our experiments we used a 1:1 molar ratio of short, dimeric DNA to dimeric TCF4. This approach led to complete dissolve of TCF4 condensates. We believe that in cellular crowded conditions, the ratios of the amounts of TRs to the DNA specific sequences are not comparable, and thus some pool of chromatin-associated TCF4 could coexist with TCF4 condensates. This suggests TCF4 IDR can form various types of interactions - hydrophobic, leading to LLPS or favorable/dominating electrostatic, with the DNA. Importantly, recent smFRET studies suggested that proteins from bHLH-LZ family, in the absence of dimerization partner form compact disordered ensembles [123]. To date, however, the oligomeric state of TCF4 in the condensate remains unknown and could be defined in the future.

**Figure 8.**
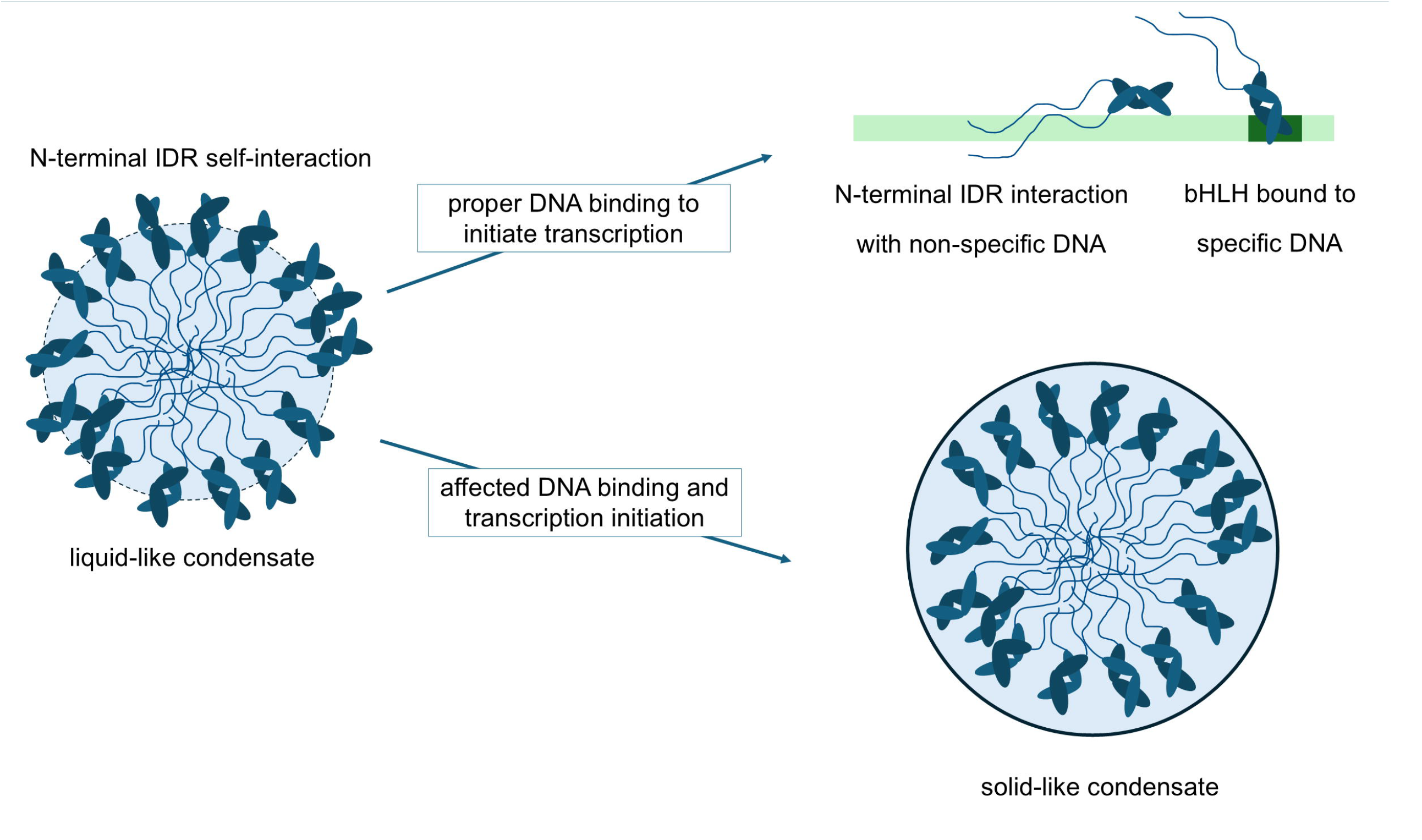

Under the conditions in which we attempted to reproduce molecular crowding using PEG 8000, TCF4 undergoes phase separation, a process that does not require elevated concentrations of NaCl and occurs in 150 mM NaCl. This process is not reversed by the addition of DNA, although short, specific DNA sequences were able to diffuse into those condensates. We take special precaution when interpreting this result. Even though PEG is one of the most used synthetic crowders and is regarded as an inert, biocompatible polymer, it has been observed that it may have an impact on protein conformation and flexibility [124]. What is more, PEG 600 molecules have been observed to form clusters through self-interaction, and those clusters were liquid-like [124,125]. Additionally, PEG 20,000 was proposed to wrap the protein, leading to the formation of aggregates [124]. Taken together, at this point it is impossible to determine whether the phase separation of TCF4 in the presence of PEG 8000 is a volume-excluded effect, or rather a side effect caused by conformational changes of TCF4 or forced clustering of TCF4 molecules due to self-association of PEG molecules interacting with TCF4.

Here we propose that the altered cellular localization pattern of some PTHS-associated TCF4 mutants may be related to LLPS. Phase separation has been previously proposed as a missing mechanism for pathogenicity of disease-linked mutations [94]. Our analysis of TCF4 lacking the C-terminal part of bHLH, substantiates the hypothesis about LLPS nature of spherical condensates present in nuclei of PTHS-linked mutants. What is more, point mutations that involve residues directly involved in interacting with DNA lead to reduced DNA binding capacity and reduced transcriptional activity of TCF4 mutants [64,89,90,120]. The result could potentially be increased TCF4 expression to offset its inefficient function. Overexpression and negative regulation of transcription mediated by TCF4 may, in turn, result in a deficiency of TCF4 molecules that leave the condensate to participate in transcription. Subsequently, TCF4 condensates can mature, and their properties change significantly. TCF4 molecules show drastic decrease in diffusion rate within condensates in the experimental time scale, despite it being a short time compared to PTHS-patient lifetime. We believe that in PTHS, TCF4 condensates can change their properties from liquid-like to solid-like, thus completely shutting down the TCF4 function. Importantly, our previous analysis suggested that PTHS-linked TCF4 point mutations may alter protein aggregation rate [126], which could be potentially enhanced by the close contact of TCF4 molecules within the condensate.

## 8. Declarations

## Ethics approval and consent to participate

Not applicable.

## Consent for publication

Not applicable.

## Availability of data and materials

The datasets generated during and/or analyzed during the current study are available from the corresponding author upon reasonable request.

## Acknowledgements

We are grateful to prof. Jerzy Dobrucki PhD, DSc. (Cell Biophysics Department, Jagiellonian University, Cracow, Poland) for cooperation and opportunity to use the Leica SP5 confocal microscope to perform CFP-TCF4/1-550, CFP-TCF4/1-601 localization imaging, and CFP-TCF4/1-601 FRAP analysis in COS-7 cells.

## Authors’ contributions

Conceptualization: A.T., Investigation: N.S., B.P.K, B.G.M, A.T., Visualization: N.S., B.P.K., A.N., B.G.M., Writing – Original Draft Preparation: N.S., B.P.K, A.N., B.G.M., A.T., Writing – Review & Editing: N.S., B.P.K., A.N., B.G.M., A.O., A.T.

## Literature

1. Banani, S.F.; Lee, H.O.; Hyman, A.A.; Rosen, M.K. Biomolecular Condensates: Organizers of Cellular Biochemistry. Nat. Rev. Mol. Cell Biol. 2017, 18, 285–298, doi:10.1038/nrm.2017.7.

2. Brangwynne, C.P.; Tompa, P.; Pappu, R. V. Polymer Physics of Intracellular Phase Transitions. Nat. Phys. 2015, 11, 899–904, doi:10.1038/nphys3532.

3. Hyman, A.A.; Weber, C.A.; Jülicher, F. Liquid-Liquid Phase Separation in Biology. Annu. Rev. Cell Dev. Biol. 2014, 30, 39–58, doi:10.1146/annurev-cellbio-100913-013325.

4. Posey, A.E.; Holehouse, A.S.; Pappu, R. V. Phase Separation of Intrinsically Disordered Proteins. Methods Enzymol. 2018, 611, 1–30, doi:10.1016/bs.mie.2018.09.035.

5. Uversky, V.N. Intrinsically Disordered Proteins in Overcrowded Milieu: Membrane-Less Organelles, Phase Separation, and Intrinsic Disorder. Curr. Opin. Struct. Biol. 2017, 44, 18–30, doi:10.1016/j.sbi.2016.10.015.

6. Mao, A.H.; Crick, S.L.; Vitalis, A.; Chicoine, C.L.; Pappu, R. V. Net Charge per Residue Modulates Conformational Ensembles of Intrinsically Disordered Proteins. Proc. Natl. Acad. Sci. 2010, 107, 8183–8188, doi:10.1073/pnas.0911107107.

7. Turoverov, K.K.; Kuznetsova, I.M.; Fonin, A. V.; Darling, A.L.; Zaslavsky, B.Y.; Uversky, V.N. Stochasticity of Biological Soft Matter: Emerging Concepts in Intrinsically Disordered Proteins and Biological Phase Separation. Trends Biochem. Sci. 2019, 44, 716–728, doi:10.1016/j.tibs.2019.03.005.

8. Darling, A.L.; Liu, Y.; Oldfield, C.J.; Uversky, V.N. Intrinsically Disordered Proteome of Human Membrane-Less Organelles. Proteomics 2018, 18, doi:10.1002/pmic.201700193.

9. Darling, A.L.; Zaslavsky, B.Y.; Uversky, V.N. Intrinsic Disorder-Based Emergence in Cellular Biology: Physiological and Pathological Liquid-Liquid Phase Transitions in Cells. Polymers (Basel). 2019, 11, 990, doi:10.3390/polym11060990.

10. Gomes, E.; Shorter, J. The Molecular Language of Membraneless Organelles. J. Biol. Chem. 2019, 294, 7115–7127, doi:10.1074/jbc.TM118.001192.

11. Shorter, J. Phasing in and Out. Nat. Chem. 2016, 8, 528–530, doi:10.1038/nchem.2534.

12. Molliex, A.; Temirov, J.; Lee, J.; Coughlin, M.; Kanagaraj, A.P.; Kim, H.J.; Mittag, T.; Taylor, J.P. Phase Separation by Low Complexity Domains Promotes Stress Granule Assembly and Drives Pathological Fibrillization. Cell 2015, 163, 123–133, doi:10.1016/j.cell.2015.09.015.

13. Mitrea, D.M.; Cika, J.A.; Guy, C.S.; Ban, D.; Banerjee, P.R.; Stanley, C.B.; Nourse, A.; Deniz, A.A.; Kriwacki, R.W. Nucleophosmin Integrates within the Nucleolus via Multi-Modal Interactions with Proteins Displaying R-Rich Linear Motifs and RRNA. Elife 2016, 5, doi:10.7554/eLife.13571.

14. Brangwynne, C.P.; Eckmann, C.R.; Courson, D.S.; Rybarska, A.; Hoege, C.; Gharakhani, J.; Jülicher, F.; Hyman, A.A. Germline P Granules Are Liquid Droplets That Localize by Controlled Dissolution/Condensation. Science (80-.). 2009, 324, 1729–1732, doi:10.1126/science.1172046.

15. Brangwynne, C.P.; Mitchison, T.J.; Hyman, A.A. Active Liquid-like Behavior of Nucleoli Determines Their Size and Shape in Xenopus Laevis Oocytes. Proc. Natl. Acad. Sci. 2011, 108, 4334–4339, doi:10.1073/pnas.1017150108.

16. Oldfield, C.J.; Dunker, A.K. Intrinsically Disordered Proteins and Intrinsically Disordered Protein Regions. Annu. Rev. Biochem. 2014, 83, 553–584, doi:10.1146/annurev-biochem-072711-164947.

17. Dyson, H.J.; Wright, P.E. Intrinsically Unstructured Proteins and Their Functions. Nat. Rev. Mol. Cell Biol. 2005, 6, 197–208, doi:10.1038/nrm1589.

18. Chong, S.; Dugast-Darzacq, C.; Liu, Z.; Dong, P.; Dailey, G.M.; Cattoglio, C.; Heckert, A.; Banala, S.; Lavis, L.; Darzacq, X.; et al. Imaging Dynamic and Selective Low-Complexity Domain Interactions That Control Gene Transcription. Science 2018, 361, eaar2555, doi:10.1126/science.aar2555.

19. Sabari, B.R.; Dall’Agnese, A.; Boija, A.; Klein, I.A.; Coffey, E.L.; Shrinivas, K.; Abraham, B.J.; Hannett, N.M.; Zamudio, A. V.; Manteiga, J.C.; et al. Coactivator Condensation at Super-Enhancers Links Phase Separation and Gene Control. Science (80-.). 2018, 361, doi:10.1126/science.aar3958.

20. Boija, A.; Klein, I.A.; Sabari, B.R.; Dall’Agnese, A.; Coffey, E.L.; Zamudio, A. V.; Li, C.H.; Shrinivas, K.; Manteiga, J.C.; Hannett, N.M.; et al. Transcription Factors Activate Genes through the Phase-Separation Capacity of Their Activation Domains. Cell 2018, 175, 1842-1855.e16, doi:10.1016/j.cell.2018.10.042.

21. Lu, H.; Yu, D.; Hansen, A.S.; Ganguly, S.; Liu, R.; Heckert, A.; Darzacq, X.; Zhou, Q. Phase-Separation Mechanism for C-Terminal Hyperphosphorylation of RNA Polymerase II. Nature 2018, 558, 318–323, doi:10.1038/s41586-018-0174-3.

22. Burke, K.A.; Janke, A.M.; Rhine, C.L.; Fawzi, N.L. Residue-by-Residue View of In Vitro FUS Granules That Bind the C-Terminal Domain of RNA Polymerase II. Mol. Cell 2015, 60, 231–241, doi:10.1016/j.molcel.2015.09.006.

23. Kwon, I.; Kato, M.; Xiang, S.; Wu, L.; Theodoropoulos, P.; Mirzaei, H.; Han, T.; Xie, S.; Corden, J.L.; McKnight, S.L. Phosphorylation-Regulated Binding of RNA Polymerase II to Fibrous Polymers of Low-Complexity Domains. Cell 2013, 155, 1049–1060, doi:10.1016/j.cell.2013.10.033.

24. Harlen, K.M.; Churchman, L.S. The Code and beyond: Transcription Regulation by the RNA Polymerase II Carboxy-Terminal Domain. Nat. Rev. Mol. Cell Biol. 2017, 18, 263–273, doi:10.1038/nrm.2017.10.

25. Cai, D.; Feliciano, D.; Dong, P.; Flores, E.; Gruebele, M.; Porat-Shliom, N.; Sukenik, S.; Liu, Z.; Lippincott-Schwartz, J. Phase Separation of YAP Reorganizes Genome Topology for Long-Term YAP Target Gene Expression. Nat. Cell Biol. 2019, 21, 1578–1589, doi:10.1038/s41556-019-0433-z.

26. Shreberk-Shaked, M.; Oren, M. New Insights into YAP/TAZ Nucleo-Cytoplasmic Shuttling: New Cancer Therapeutic Opportunities? Mol. Oncol. 2019, 13, 1335–1341, doi:10.1002/1878-0261.12498.

27. Lu, Y.; Wu, T.; Gutman, O.; Lu, H.; Zhou, Q.; Henis, Y.I.; Luo, K. Phase Separation of TAZ Compartmentalizes the Transcription Machinery to Promote Gene Expression. Nat. Cell Biol. 2020, 22, 453–464, doi:10.1038/s41556-020-0485-0.

28. Strom, A.R.; Emelyanov, A. V; Mir, M.; Fyodorov, D. V; Darzacq, X.; Karpen, G.H. Phase Separation Drives Heterochromatin Domain Formation. Nature 2017, 547, 241–245, doi:10.1038/nature22989.

29. Shin, Y.; Chang, Y.-C.; Lee, D.S.W.; Berry, J.; Sanders, D.W.; Ronceray, P.; Wingreen, N.S.; Haataja, M.; Brangwynne, C.P. Liquid Nuclear Condensates Mechanically Sense and Restructure the Genome. Cell 2018, 175, 1481-1491.e13, doi:10.1016/j.cell.2018.10.057.

30. Quinodoz, S.A.; Ollikainen, N.; Tabak, B.; Palla, A.; Schmidt, J.M.; Detmar, E.; Lai, M.M.; Shishkin, A.A.; Bhat, P.; Takei, Y.; et al. Higher-Order Inter-Chromosomal Hubs Shape 3D Genome Organization in the Nucleus. Cell 2018, 174, 744-757.e24, doi:10.1016/j.cell.2018.05.024.

31. Gibson, B.A.; Doolittle, L.K.; Schneider, M.W.G.; Jensen, L.E.; Gamarra, N.; Henry, L.; Gerlich, D.W.; Redding, S.; Rosen, M.K. Organization of Chromatin by Intrinsic and Regulated Phase Separation. Cell 2019, 179, 470-484.e21, doi:10.1016/j.cell.2019.08.037.

32. Larson, A.G.; Elnatan, D.; Keenen, M.M.; Trnka, M.J.; Johnston, J.B.; Burlingame, A.L.; Agard, D.A.; Redding, S.; Narlikar, G.J. Liquid Droplet Formation by HP1α Suggests a Role for Phase Separation in Heterochromatin. Nature 2017, 547, 236–240, doi:10.1038/nature22822.

33. Narlikar, G.J. Phase-Separation in Chromatin Organization. J. Biosci. 2020, 45.

34. Langdon, E.M.; Gladfelter, A.S. A New Lens for RNA Localization: Liquid-Liquid Phase Separation. Annu. Rev. Microbiol. 2018, 72, 255–271, doi:10.1146/annurev-micro-090817-062814.

35. Hu, J.; Khodadadi-Jamayran, A.; Mao, M.; Shah, K.; Yang, Z.; Nasim, M.T.; Wang, Z.; Jiang, H. AKAP95 Regulates Splicing through Scaffolding RNAs and RNA Processing Factors. Nat. Commun. 2016, 7, 13347, doi:10.1038/ncomms13347.

36. Li, W.; Hu, J.; Shi, B.; Palomba, F.; Digman, M.A.; Gratton, E.; Jiang, H. Biophysical Properties of AKAP95 Protein Condensates Regulate Splicing and Tumorigenesis. Nat. Cell Biol. 2020, 22, 960–972, doi:10.1038/s41556-020-0550-8.

37. Xing, W.; Muhlrad, D.; Parker, R.; Rosen, M.K. A Quantitative Inventory of Yeast P Body Proteins Reveals Principles of Composition and Specificity. Elife 2020, 9, doi:10.7554/eLife.56525.

38. Lécuyer, E.; Yoshida, H.; Parthasarathy, N.; Alm, C.; Babak, T.; Cerovina, T.; Hughes, T.R.; Tomancak, P.; Krause, H.M. Global Analysis of MRNA Localization Reveals a Prominent Role in Organizing Cellular Architecture and Function. Cell 2007, 131, 174–187, doi:10.1016/j.cell.2007.08.003.

39. Mirza-Aghazadeh-Attari, M.; Mohammadzadeh, A.; Yousefi, B.; Mihanfar, A.; Karimian, A.; Majidinia, M. 53BP1: A Key Player of DNA Damage Response with Critical Functions in Cancer. DNA Repair (Amst). 2019, 73, 110–119, doi:10.1016/j.dnarep.2018.11.008.

40. Pessina, F.; Giavazzi, F.; Yin, Y.; Gioia, U.; Vitelli, V.; Galbiati, A.; Barozzi, S.; Garre, M.; Oldani, A.; Flaus, A.; et al. Functional Transcription Promoters at DNA Double-Strand Breaks Mediate RNA-Driven Phase Separation of Damage-Response Factors. Nat. Cell Biol. 2019, 21, 1286– 1299, doi:10.1038/s41556-019-0392-4.

41. Kilic, S.; Lezaja, A.; Gatti, M.; Bianco, E.; Michelena, J.; Imhof, R.; Altmeyer, M. Phase Separation of 53BP1 Determines Liquid-like Behavior of DNA Repair Compartments. EMBO J. 2019, 38, e101379, doi:10.15252/embj.2018101379.

42. Teloni, F.; Altmeyer, M. Readers of Poly(ADP-Ribose): Designed to Be Fit for Purpose. Nucleic Acids Res. 2016, 44, 993–1006, doi:10.1093/nar/gkv1383.

43. Singatulina, A.S.; Hamon, L.; Sukhanova, M. V; Desforges, B.; Joshi, V.; Bouhss, A.; Lavrik, O.I.; Pastré, D. PARP-1 Activation Directs FUS to DNA Damage Sites to Form PARG-Reversible Compartments Enriched in Damaged DNA. Cell Rep. 2019, 27, 1809-1821.e5, doi:10.1016/j.celrep.2019.04.031.

44. Cuella-Martin, R.; Oliveira, C.; Lockstone, H.E.; Snellenberg, S.; Grolmusova, N.; Chapman, J.R. 53BP1 Integrates DNA Repair and P53-Dependent Cell Fate Decisions via Distinct Mechanisms. Mol. Cell 2016, 64, 51–64, doi:10.1016/j.molcel.2016.08.002.

45. Cai, D.; Liu, Z.; Lippincott-Schwartz, J. Biomolecular Condensates and Their Links to Cancer Progression. Trends Biochem. Sci. 2021, 46, 535–549, doi:10.1016/j.tibs.2021.01.002.

46. Galloux, M.; Longhi, S. Unraveling Liquid–Liquid Phase Separation (LLPS) in Viral Infections to Understand and Treat Viral Diseases. Int. J. Mol. Sci. 2024, 25, 6981, doi:10.3390/ijms25136981.

47. Wang, L.; Wang, Y.; Ke, Z.; Wang, Z.; Guo, Y.; Zhang, Y.; Zhang, X.; Guo, Z.; Wan, B. Liquid-Liquid Phase Separation: A New Perspective on Respiratory Diseases. Front. Immunol. 2024, 15, 1444253, doi:10.3389/fimmu.2024.1444253.

48. Zhang, X.; Yuan, L.; Zhang, W.; Zhang, Y.; Wu, Q.; Li, C.; Wu, M.; Huang, Y. Liquid-Liquid Phase Separation in Diseases. MedComm 2024, 5, e640, doi:10.1002/mco2.640.

49. Wu, Y.; Ma, B.; Liu, C.; Li, D.; Sui, G. Pathological Involvement of Protein Phase Separation and Aggregation in Neurodegenerative Diseases. Int. J. Mol. Sci. 2024, 25, 10187, doi:10.3390/ijms251810187.

50. Liu, Z.; Qin, Z.; Liu, Y.; Xia, X.; He, L.; Chen, N.; Hu, X.; Peng, X. Liquid1liquid Phase Separation: Roles and Implications in Future Cancer Treatment. Int. J. Biol. Sci. 2023, 19, 4139–4156, doi:10.7150/ijbs.81521.

51. Wang, B.; Zhang, L.; Dai, T.; Qin, Z.; Lu, H.; Zhang, L.; Zhou, F. Liquid–Liquid Phase Separation in Human Health and Diseases. Signal Transduct. Target. Ther. 2021, 6, 290, doi:10.1038/s41392-021-00678-1.

52. Zbinden, A.; Pérez-Berlanga, M.; De Rossi, P.; Polymenidou, M. Phase Separation and Neurodegenerative Diseases: A Disturbance in the Force. Dev. Cell 2020, 55, 45–68, doi:10.1016/j.devcel.2020.09.014.

53. Massari, M.E.; Murre, C. Helix-Loop-Helix Proteins: Regulators of Transcription in Eucaryotic Organisms. Mol. Cell. Biol. 2000, 20, 429–440, doi:10.1128/MCB.20.2.429-440.2000.

54. Murre, C.; McCaw, P.S.; Baltimore, D. A New DNA Binding and Dimerization Motif in Immunoglobulin Enhancer Binding, Daughterless, MyoD, and Myc Proteins. Cell 1989, 56, 777–783, doi:10.1016/0092-8674(89)90682-X.

55. Ellenberger, T.; Arnaud, M.; Harrison, S.C. Crystal Structure of Transcription Factor E471: E-Box Recognition by a Basic Region Helix-Loop-Helix Dimer. Genes Dev. 1994, 8, 970–980, doi:10.1101/gad.8.8.970.

56. Voronova, Anna; Baltimore, D. Mutations That Disrupt DNA Binding and Dimer Formation in the E47 Helix-Loop-Helix Protein Map to Distinct Domains. Proc. Natl. Acad. Sci. U. S. A. 1990, 87, 4722–4726, doi:10.1073/pnas.87.12.4722.

57. Murre, C.; McCaw, P.S.; Vaessin, H.; Caudy, M.; Jan, L.Y.; Jan, Y.N.; Cabrera, C. V.; Buskin, J.N.; Hauschka, S.D.; Lassar, A.B.; et al. Interactions between Heterologous Helix-Loop-Helix Proteins Generate Complexes That Bind Specifically to a Common DNA Sequence. Cell 1989, 58, 537–544, doi:10.1016/0092-8674(89)90434-0.

58. Henthorn, P.; Kiledjian, M.; Kadesch, T. Two Distinct Transcription Factors That Bind the Immunoglobulin Enhancer ?E5/?E2 Motif. Science (80-.). 1990, 247, 467–470, doi:10.1126/science.2105528.

59. Corneliussen, B.; Thornell, A.; Hallberg, B.; Grundström, T. Helix-Loop-Helix Transcriptional Activators Bind to a Sequence in Glucocorticoid Response Elements of Retrovirus Enhancers. J. Virol. 1991, 65, 6084–6093, doi:10.1128/jvi.65.11.6084-6093.1991.

60. Pscherer, A.; Dörflinger, U.; Kirfel, J.; Gawlas, K.; Rüschoff, J.; Buettner, R.; Schüle, R. The Helix-Loop-Helix Transcription Factor SEF-2 Regulates the Activity of a Novel Initiator Element in the Promoter of the Human Somatostatin Receptor II Gene. EMBO J. 1996, 15, 6680–6690, doi:10.1002/j.1460-2075.1996.tb01058.x.

61. Yoon, S.O.; Chikaraishi, D.M. Isolation of Two E-Box Binding Factors That Interact with the Rat Tyrosine Hydroxylase Enhancer. J. Biol. Chem. 1994, 269, 18453–18462, doi:10.1016/S0021-9258(17)32330-X.

62. Longo, A.; Guanga, G.P.; Rose, R.B. Crystal Structure of E47-NeuroD1/Beta2 BHLH Domain-DNA Complex: Heterodimer Selectivity and DNA Recognition. Biochemistry 2008, 47, 218–229, doi:10.1021/bi701527r.

63. De Pontual, L.; Mathieu, Y.; Golzio, C.; Rio, M.; Malan, V.; Boddaert, N.; Soufflet, C.; Picard, C.; Durandy, A.; Dobbie, A.; et al. Mutational, Functional, and Expression Studies of the TCF4 Gene in Pitt-Hopkins Syndrome. Hum. Mutat. 2009, 30, 669–676, doi:10.1002/humu.20935.

64. Forrest, M.; Chapman, R.M.; Doyle, A.M.; Tinsley, C.L.; Waite, A.; Blake, D.J. Functional Analysis of TCF4 Missense Mutations That Cause Pitt-Hopkins Syndrome. Hum. Mutat. 2012, 33, 1676–1686, doi:10.1002/humu.22160.

65. Petropoulos, H.; Skerjanc, I.S. Analysis of the Inhibition of MyoD Activity by ITF-2B and Full-Length E12/E47. J. Biol. Chem. 2000, 275, 25095–25101, doi:10.1074/jbc.M004251200.

66. Lu, Y.; Sheng, D.-Q.; Mo, Z.-C.; Li, H.-F.; Wu, N.-H.; Shen, Y.-F. A Negative Regulatory Element-Dependent Inhibitory Role of ITF2B on IL-2 Receptor α Gene. Biochem. Biophys. Res. Commun. 2005, 336, 142–149, doi:10.1016/j.bbrc.2005.08.050.

67. Furumura, M.; Potterf, S.B.; Toyofuku, K.; Matsunaga, J.; Muller, J.; Hearing, V.J. Involvement of ITF2 in the Transcriptional Regulation of Melanogenic Genes. J. Biol. Chem. 2001, 276, 28147–28154, doi:10.1074/jbc.M101626200.

68. Sepp, M.; Kannike, K.; Eesmaa, A.; Urb, M. Functional Diversity of Human Basic Helix-Loop-Helix Transcription Factor TCF4 Isoforms Generated by Alternative 5 9 Exon Usage and Splicing. PLoS One 2011, 6, doi:10.1371/journal.pone.0022138.

69. Aronheim, A.; Shiran, R.; Rosen, A.; Walker, M.D. The E2A Gene Product Contains Two Separable and Functionally Distinct Transcription Activation Domains. Proc. Natl. Acad. Sci. U. S. A. 1993, doi:10.1073/pnas.90.17.8063.

70. Quong, M.W.; Massari, M.E.; Zwart, R.; Murre, C. A New Transcriptional-Activation Motif Restricted to a Class of Helix-Loop-Helix Proteins Is Functionally Conserved in Both Yeast and Mammalian Cells. Mol. Cell. Biol. 1993, 13, 792–800, doi:10.1128/mcb.13.2.792.

71. Massari, M.E.; Jennings, P.A.; Murre, C. The AD1 Transactivation Domain of E2A Contains a Highly Conserved Helix Which Is Required for Its Activity in Both Saccharomyces Cerevisiae and Mammalian Cells. Mol. Cell. Biol. 1996, 16, 121–129, doi:10.1128/mcb.16.1.121.

72. Bayly, R.; Chuen, L.; Currie, R.A.; Hyndman, B.D.; Casselman, R.; Blobel, G.A.; LeBrun, D.P. E2A-PBX1 Interacts Directly with the KIX Domain of CBP/P300 in the Induction of Proliferation in Primary Hematopoietic Cells. J. Biol. Chem. 2004, doi:10.1074/jbc.M408654200.

73. Denis, C.M.; Chitayat, S.; Plevin, M.J.; Wang, F.; Thompson, P.; Liu, S.; Spencer, H.L.; Ikura, M.; LeBrun, D.P.; Smith, S.P. Structural Basis of CBP/P300 Recruitment in Leukemia Induction by E2A-PBX1. Blood 2012, doi:10.1182/blood-2012-02-411397.

74. Denis, C.M.; Langelaan, D.N.; Kirlin, A.C.; Chitayat, S.; Munro, K.; Spencer, H.L.; Lebrun, D.P.; Smith, S.P. Functional Redundancy between the Transcriptional Activation Domains of E2A Is Mediated by Binding to the KIX Domain of CBP/P300. Nucleic Acids Res. 2014, doi:10.1093/nar/gku206.

75. Massari, M.E.; Grant, P.A.; Pray-Grant, M.G.; Berger, S.L.; Workman, J.L.; Murre, C. A Conserved Motif Present in a Class of Helix-Loop-Helix Proteins Activates Transcription by Direct Recruitment of the SAGA Complex. Mol. Cell 1999, doi:10.1016/S1097-2765(00)80188-4.

76. Zhang, J.; Kalkum, M.; Yamamura, S.; Chait, B.T.; Roeder, R.G. E Protein Silencing by the Leukemogenic AML1-ETO Fusion Protein. Science (80-.). 2004, doi:10.1126/science.1097937.

77. Herbst, A.; Kolligs, F.T. A Conserved Domain in the Transcription Factor ITF-2B Attenuates Its Activity. Biochem. Biophys. Res. Commun. 2008, doi:10.1016/j.bbrc.2008.03.081.

78. Markus, M.; Du, Z.; Benezra, R. Enhancer-Specific Modulation of E Protein Activity. J. Biol. Chem. 2002, doi:10.1074/jbc.M110659200.

79. Greb-Markiewicz, B.; Kazana, W.; Zarębski, M.; Ożyhar, A. The Subcellular Localization of BHLH Transcription Factor TCF4 Is Mediated by Multiple Nuclear Localization and Nuclear Export Signals. Sci. Rep. 2019, 9, 15629, doi:10.1038/s41598-019-52239-w.

80. D’Rozario, M.; Zhang, T.; Waddell, E.A.; Zhang, Y.; Sahin, C.; Sharoni, M.; Hu, T.; Nayal, M.; Kutty, K.; Liebl, F.; et al. Type I BHLH Proteins Daughterless and Tcf4 Restrict Neurite Branching and Synapse Formation by Repressing Neurexin in Postmitotic Neurons. Cell Rep. 2016, 15, 386–397, doi:10.1016/j.celrep.2016.03.034.

81. Sozańska, N.; Klepka, B.P.; Niedzwiecka, A.; Zhukova, L.; Dadlez, M.; Greb-Markiewicz, B.; Ożyhar, A.; Tarczewska, A. The Molecular Properties of the BHLH TCF4 Protein as an Intrinsically Disordered Hub Transcription Factor. Cell Commun. Signal. 2025, 23, 154, doi:10.1186/s12964-025-02154-7.

82. Quednow, B.B.; Brzózka, M.M.; Rossner, M.J. Transcription Factor 4 (TCF4) and Schizophrenia: Integrating the Animal and the Human Perspective. Cell. Mol. Life Sci. 2014, 71, 2815–2835, doi:10.1007/s00018-013-1553-4.

83. Forrest, M.P.; Hill, M.J.; Quantock, A.J.; Martin-Rendon, E.; Blake, D.J. The Emerging Roles of TCF4 in Disease and Development. Trends Mol. Med. 2014, 20, 322–331, doi:10.1016/j.molmed.2014.01.010.

84. Pitt, D.; Hopkins, I. A Syndrome of Mental Retardation, Wide Mouth and Intermittent Overbreathing. Aust. Paediatr. J. 1978, doi:10.1111/jpc.1978.14.3.182.

85. Sweetser, D.A.; Elsharkawi, I.; Yonker, L.; Steeves, M.; Parkin, K.; Thibert, R. Pitt-Hopkins Syndrome; Adam, M.P., Feldman, J., Mirzaa, G.M., Pagon, R.A., Wallace, S.E., Amemiya, A., Eds.; 2018;

86. Amiel, J.; Rio, M.; Pontual, L. de; Redon, R.; Malan, V.; Boddaert, N.; Plouin, P.; Carter, N.P.; Lyonnet, S.; Munnich, A.; et al. Mutations in TCF4, Encoding a Class I Basic Helix-Loop-Helix Transcription Factor, Are Responsible for Pitt-Hopkins Syndrome, a Severe Epileptic Encephalopathy Associated with Autonomic Dysfunction. Am. J. Hum. Genet. 2007, 80, 988– 993, doi:10.1086/515582.

87. Zweier, C.; Peippo, M.M.; Hoyer, J.; Sousa, S.; Bottani, A.; Clayton-Smith, J.; Reardon, W.; Saraiva, J.; Cabral, A.; Göhring, I.; et al. Haploinsufficiency of TCF4 Causes Syndromal Mental Retardation with Intermittent Hyperventilation (Pitt-Hopkins Syndrome). Am. J. Hum. Genet. 2007, 80, 994–1001, doi:10.1086/515583.

88. Brockschmidt, A.; Todt, U.; Ryu, S.; Hoischen, A.; Landwehr, C.; Birnbaum, S.; Frenck, W.; Radlwimmer, B.; Lichter, P.; Engels, H.; et al. Severe Mental Retardation with Breathing Abnormalities (Pitt - Hopkins Syndrome) Is Caused by Haploinsufficiency of the Neuronal BHLH Transcription Factor TCF4. Hum. Mol. Genet. 2007, 16, 1488–1494, doi:10.1093/hmg/ddm099.

89. Sepp, M.; Pruunsild, P.; Timmusk, T. Pitt–Hopkins Syndrome-Associated Mutations in TCF4 Lead to Variable Impairment of the Transcription Factor Function Ranging from Hypomorphic to Dominant-Negative Effects. Hum. Mol. Genet. 2012, 21, 2873–2888, doi:10.1093/hmg/dds112.

90. Sirp, A.; Roots, K.; Nurm, K.; Tuvikene, J.; Sepp, M.; Timmusk, T. Functional Consequences of TCF4 Missense Substitutions Associated with Pitt-Hopkins Syndrome, Mild Intellectual Disability, and Schizophrenia. J. Biol. Chem. 2021, 297, 101381, doi:10.1016/j.jbc.2021.101381.

91. Stefansson, H.; Ophoff, R.A.; Steinberg, S.; Andreassen, O.A.; Cichon, S.; Rujescu, D.; Werge, T.; Pietiläinen, O.P.H.; Mors, O.; Mortensen, P.B.; et al. Common Variants Conferring Risk of Schizophrenia. Nature 2009, 460, 744–747, doi:10.1038/nature08186.

92. Hu, X.; Zhang, B.; Liu, W.; Paciga, S.; He, W.; Lanz, T.A.; Kleiman, R.; Dougherty, B.; Hall, S.K.; McIntosh, A.M.; et al. A Survey of Rare Coding Variants in Candidate Genes in Schizophrenia by Deep Sequencing. Mol. Psychiatry 2014, 19, 858–859, doi:10.1038/mp.2013.131.

93. Basmanav, F.B.; Forstner, A.J.; Fier, H.; Herms, S.; Meier, S.; Degenhardt, F.; Hoffmann, P.; Barth, S.; Fricker, N.; Strohmaier, J.; et al. Investigation of the Role of TCF4 Rare Sequence Variants in Schizophrenia. Am. J. Med. Genet. Part B Neuropsychiatr. Genet. 2015, 168, 354– 362, doi:10.1002/ajmg.b.32318.

94. Tsang, B.; Pritišanac, I.; Scherer, S.W.; Moses, A.M.; Forman-Kay, J.D. Phase Separation as a Missing Mechanism for Interpretation of Disease Mutations. Cell 2020, 183, 1742–1756, doi:10.1016/j.cell.2020.11.050.

95. Hardenberg, M.; Horvath, A.; Ambrus, V.; Fuxreiter, M.; Vendruscolo, M. Widespread Occurrence of the Droplet State of Proteins in the Human Proteome. Proc. Natl. Acad. Sci. U. S. A. 2020, 117, 33254–33262, doi:10.1073/PNAS.2007670117.

96. Vendruscolo, M.; Fuxreiter, M. Sequence Determinants of the Aggregation of Proteins Within Condensates Generated by Liquid-Liquid Phase Separation. J. Mol. Biol. 2022, 434, 167201, doi:10.1016/j.jmb.2021.167201.

97. Hatos, A.; Tosatto, S.C.E.; Vendruscolo, M.; Fuxreiter, M. FuzDrop on AlphaFold: Visualizing the Sequence-Dependent Propensity of Liquid–Liquid Phase Separation and Aggregation of Proteins. Nucleic Acids Res. 2022, 50, W337–W344, doi:10.1093/nar/gkac386.

98. Vernon, R.M.C.; Chong, P.A.; Tsang, B.; Kim, T.H.; Bah, A.; Farber, P.; Lin, H.; Forman-Kay, J.D. Pi-Pi Contacts Are an Overlooked Protein Feature Relevant to Phase Separation. Elife 2018, 7, 1–48, doi:10.7554/eLife.31486.

99. Ibrahim, A.Y.; Khaodeuanepheng, N.P.; Amarasekara, D.L.; Correia, J.J.; Lewis, K.A.; Fitzkee, N.C.; Hough, L.E.; Whitten, S.T. Intrinsically Disordered Regions That Drive Phase Separation Form a Robustly Distinct Protein Class. J. Biol. Chem. 2023, 299, 102801, doi:10.1016/j.jbc.2022.102801.

100. Bolognesi, B.; Gotor, N.L.; Dhar, R.; Cirillo, D.; Baldrighi, M.; Tartaglia, G.G.; Lehner, B. A Concentration-Dependent Liquid Phase Separation Can Cause Toxicity upon Increased Protein Expression. Cell Rep. 2016, 16, 222–231, doi:10.1016/j.celrep.2016.05.076.

101. Fluorescence Polarization. Emission Anisotropy. In Molecular Fluorescence; Wiley, 2001; pp. 125–154.

102. Kroschwald, S.; Maharana, S.; Simon, A. Hexanediol: A Chemical Probe to Investigate the Material Properties of Membrane-Less Compartments. Matters 2017, doi:10.19185/matters.201702000010.

103. Feng, Z.; Chen, X.; Wu, X.; Zhang, M. Formation of Biological Condensates via Phase Separation: Characteristics, Analytical Methods, and Physiological Implications. J. Biol. Chem. 2019, 294, 14823–14835, doi:10.1074/jbc.REV119.007895.

104. Yang, J.; Horton, J.R.; Li, J.; Huang, Y.; Zhang, X.; Blumenthal, R.M.; Cheng, X. Structural Basis for Preferential Binding of Human TCF4 to DNA Containing 5-Carboxylcytosine. Nucleic Acids Res. 2019, 47, 8375–8387, doi:10.1093/nar/gkz381.

105. Wright, P.E.; Dyson, H.J. Intrinsically Disordered Proteins in Cellular Signalling and Regulation. Nat. Rev. Mol. Cell Biol. 2015, 16, 18–29, doi:10.1038/nrm3920.

106. Batlle, C.; Yang, P.; Coughlin, M.; Messing, J.; Pesarrodona, M.; Szulc, E.; Salvatella, X.; Kim, H.J.; Taylor, J.P.; Ventura, S. HnRNPDL Phase Separation Is Regulated by Alternative Splicing and Disease-Causing Mutations Accelerate Its Aggregation. Cell Rep. 2020, 30, 1117-1128.e5, doi:10.1016/j.celrep.2019.12.080.

107. McNally, J.G.; Müller, W.G.; Walker, D.; Wolford, R.; Hager, G.L. The Glucocorticoid Receptor: Rapid Exchange with Regulatory Sites in Living Cells. Science (80-.). 2000, 287, 1262–1265, doi:10.1126/science.287.5456.1262.

108. Elf, J.; Li, G.-W.; Xie, X.S. Probing Transcription Factor Dynamics at the Single-Molecule Level in a Living Cell. Science (80-.). 2007, 316, 1191–1194, doi:10.1126/science.1141967.

109. de Jonge, W.J.; Patel, H.P.; Meeussen, J.V.W.; Lenstra, T.L. Following the Tracks: How Transcription Factor Binding Dynamics Control Transcription. Biophys. J. 2022, 121, 1583– 1592, doi:10.1016/j.bpj.2022.03.026.

110. Mazzocca, M.; Fillot, T.; Loffreda, A.; Gnani, D.; Mazza, D. The Needle and the Haystack: Single Molecule Tracking to Probe the Transcription Factor Search in Eukaryotes. Biochem. Soc. Trans. 2021, 49, 1121–1132, doi:10.1042/BST20200709.

111. Suter, D.M. Transcription Factors and DNA Play Hide and Seek. Trends Cell Biol. 2020, 30, 491– 500, doi:10.1016/j.tcb.2020.03.003.

112. Chen, J.; Zhang, Z.; Li, L.; Chen, B.-C.; Revyakin, A.; Hajj, B.; Legant, W.; Dahan, M.; Lionnet, T.; Betzig, E.; et al. Single-Molecule Dynamics of Enhanceosome Assembly in Embryonic Stem Cells. Cell 2014, 156, 1274–1285, doi:10.1016/j.cell.2014.01.062.

113. Morisaki, T.; Müller, W.G.; Golob, N.; Mazza, D.; McNally, J.G. Single-Molecule Analysis of Transcription Factor Binding at Transcription Sites in Live Cells. Nat. Commun. 2014, 5, 4456, doi:10.1038/ncomms5456.

114. Lu, F.; Lionnet, T. Transcription Factor Dynamics. Cold Spring Harb. Perspect. Biol. 2021, 13, a040949, doi:10.1101/cshperspect.a040949.

115. Pabo, C.O.; Sauer, R.T. TRANSCRIPTION FACTORS: Structural Families and Principles of DNA Recognition. Annu. Rev. Biochem. 1992, 61, 1053–1095, doi:10.1146/annurev.bi.61.070192.005201.

116. Callegari, A.; Sieben, C.; Benke, A.; Suter, D.M.; Fierz, B.; Mazza, D.; Manley, S. Single-Molecule Dynamics and Genome-Wide Transcriptomics Reveal That NF-KB (P65)-DNA Binding Times Can Be Decoupled from Transcriptional Activation. PLOS Genet. 2019, 15, e1007891, doi:10.1371/journal.pgen.1007891.

117. Shao, S.; Zhang, H.; Zeng, Y.; Li, Y.; Sun, C.; Sun, Y. TagBiFC Technique Allows Long-Term Single-Molecule Tracking of Protein-Protein Interactions in Living Cells. Commun. Biol. 2021, 4, 378, doi:10.1038/s42003-021-01896-7.

118. Xie, L.; Torigoe, S.E.; Xiao, J.; Mai, D.H.; Li, L.; Davis, F.P.; Dong, P.; Marie-Nelly, H.; Grimm, J.; Lavis, L.; et al. A Dynamic Interplay of Enhancer Elements Regulates Klf4 Expression in Naïve Pluripotency. Genes Dev. 2017, 31, 1795–1808, doi:10.1101/gad.303321.117.

119. Shin, Y.; Brangwynne, C.P. Liquid Phase Condensation in Cell Physiology and Disease. Science (80-.). 2017, 357, doi:10.1126/science.aaf4382.

120. Zweier, C.; Sticht, H.; Bijlsma, E.K.; Clayton-Smith, J.; Boonen, S.E.; Fryer, A.; Greally, M.T.; Hoffmann, L.; den Hollander, N.S.; Jongmans, M.; et al. Further Delineation of Pitt-Hopkins Syndrome: Phenotypic and Genotypic Description of 16 Novel Patients. J. Med. Genet. 2008, 45, 738–744, doi:10.1136/jmg.2008.060129.

121. Giurgea, I.; Missirian, C.; Cacciagli, P.; Whalen, S.; Fredriksen, T.; Gaillon, T.; Rankin, J.; Mathieu-Dramard, M.; Morin, G.; Martin-Coignard, D.; et al. TCF4 Deletions in Pitt-Hopkins Syndrome. Hum. Mutat. 2008, 29, 242–251, doi:10.1002/humu.20859.

122. Whalen, S.; Héron, D.; Gaillon, T.; Moldovan, O.; Rossi, M.; Devillard, F.; Giuliano, F.; Soares, G.; Mathieu-Dramard, M.; Afenjar, A.; et al. Novel Comprehensive Diagnostic Strategy in Pitt-Hopkins Syndrome: Clinical Score and Further Delineation of the TCF4 Mutational Spectrum. Hum. Mutat. 2012, 33, 64–72, doi:10.1002/humu.21639.

123. Vancraenenbroeck, R.; Harel, Y.S.; Zheng, W.; Hofmann, H. Polymer Effects Modulate Binding Affinities in Disordered Proteins. Proc. Natl. Acad. Sci. 2019, 116, 19506–19512, doi:10.1073/pnas.1904997116.

124. Liebau, J.; Laatsch, B.F.; Rusnak, J.; Gunderson, K.; Finke, B.; Bargender, K.; Narkiewicz-Jodko, A.; Weeks, K.; Williams, M.T.; Shulgina, I.; et al. Polyethylene Glycol Impacts Conformation and Dynamics of Escherichia Coli Prolyl-TRNA Synthetase Via Crowding and Confinement Effects. Biochemistry 2024, 63, 1621–1635, doi:10.1021/acs.biochem.3c00719.

125. Laatsch, B.F.; Brandt, M.; Finke, B.; Fossum, C.J.; Wackett, M.J.; Lowater, H.R.; Narkiewicz-Jodko, A.; Le, C.N.; Yang, T.; Glogowski, E.M.; et al. Polyethylene Glycol 20k. Does It Fluoresce? ACS Omega 2023, 8, 14208–14218, doi:10.1021/acsomega.3c01124.

126. Sozańska, N.; Krupnik, V.; Greb-Markiewicz, B.; Ożyhar, A.; Tarczewska, A. An Attempt to Explain What Intrinsically Disordered TCF4 Does in Its Spare Time When PTHS-Related Mutations Prevent It from Doing Its Job. Cell Commun. Signal. 2025, 23, 258, doi:10.1186/s12964-025-02265-1.

