## Supplementary material for "Intrinsically disordered bHLH family member TCF4 drives LLPS in a DNA-dependent manner": captions

**Fig. 1** (A) Subcellular distribution of CFP tagged deletion mutants of TCF4-B^+^ isoform. Representative images (single confocal plane for confocal microscopy) of expressed CFP-TCF4-B^+^/1-550 and CFP-TCF4-B^+^/1-601 in COS-7 cells. Subcellular localizations of the expressed proteins were analyzed by confocal microscopy 24 h after COS-7 cells transfection. Scale bar is 10 µm. (B) FRAP verification of liquid character of CFP-TCF4-B^+^/1-601 expressed in COS-7 cells, 24 after transfection. Scale bar is 5 µm. (C) The magnification of ROI in the FRAP experiment presented in C. Scale bar is 0.5 µm. (D) Addition of 10% 1,6-hexanediol results in dissolving condensates of the expressed CFP-TCF4-B^+^/1-601 in COS-7 cells in less than 2 s. Scale bar is 10 µm.

**Fig. 2** Propensity of TCF4 to LLPS predicted by (A) FuzDrop, (B) PScore, (C) ParSe, and (D) catGRANULE. (A-D) gray area corresponds to bHLH, the only folded domain of TCF4. (A, B, D) Black dashed line represents threshold and values above it indicate high probability that the protein fragments can drive LLPS. (C) P (blue) are the regions predicted as intrinsically disordered and prone to undergo phase separation; D (red) are classified as intrinsically disordered but do not undergo phase separation; F regions (black) may or may not be intrinsically disordered, but can fold to a stable conformation. (E) Fluorescence confocal microscopy images of coalescence of condensates formed in solution of TCF4 at 1 mg/mL in a buffer containing 500 mM NaCl. Recording of the movie (Supplemental Video) started after 3 min of incubation. Scale bar is the same for all images.

**Fig. 3** (A) Fluorescence confocal microscopy images of condensates formed in solution of TCF4 at 1 mg/mL, at increasing NaCl concentrations, after 3 and 5 min of incubation. (B) Fluorescence recovery after photobleaching images shown for the lowest and highest NaCl concentrations measured. FRAP analysis as a function of salt concentration performed after (C) 3 and (D) 5 min of incubation. Dependence of (E) mobile fraction and (F) recovery half-time on NaCl concentration. Data points shown with SD; lines, fitted functions; shaded area, 95% confidence interval. In A and B, the scale bar is the same for all images.

**Fig. 4** (A) Fluorescence confocal microscopy images showing the time evolution of liquid phases formed in solution of TCF4 at 1 mg/mL, at 500 mM NaCl. (B) FRAP images shown for different incubation times. (C) FRAP analysis as a function of incubation time at 500 mM NaCl. (D) Mobile fraction and recovery half-time as a function of incubation time. Data points shown with SD; lines, fitted functions; shaded area, 95% confidence interval. In A and B, the scale bar is the same for all images.

**Fig. 5** (A) Phase diagram determined by widefield microscopy for TCF4. Colored areas are visual aids representing homogenous solution (white), droplets (blue), and different types of assemblies (gray) present in the sample analyzed. (B-D) Representative DIC microscopic images of TCF4 samples at different NaCl concentrations. Scale bar is 10 μM and is the same for all images. TCF4 concentration is 1.5 mg/mL and NaCl concentrations are (B) 400 mM, (C) 600 mM, (D) 900 mM. (E-G) Representative fluorescent microscopic images of TCF4 samples at different NaCl concentrations. TCF4 concentration is 1.5 mg/mL and NaCl concentrations are (E) 800 mM, (F) 1 M, (G) 1.5 M.

**Fig. 6** DIC microscopic images of TCF4 samples (A) after addition of: 20 μM dsE-box, 20 μM ssE-box, 20 μM non-specific dsDNA, or (B) buffer S. The concentrations were 1 mg/mL TCF4 and 700 mM NaCl in all the samples. Scale bar is 10 μM and is the same for all images.

**Fig. 7** Determination of the ability of TCF4 to undergo LLPS under molecular crowding conditions. (A-E) Fluorescent microscopic images of TCF4 (including ∼5% of fluorescently labeled protein) at 0.5 mg/mL, in buffer S in (A) the absence of PEG 8000 and presence of PEG 8000 at (B) 3%, (C) 5%, (D) 7%, (E) 10%. The scale bar is 5 μm, the same for all images. (F) Dependence of TCF4 fluorescence anisotropy on PEG 8000 concentration after 5 min of PEG addition. (G) DIC microscopic image of unlabeled TCF4 at 0.5 mg/mL, in buffer S in the presence of 5% PEG 8000. (H) Fluorescent microscopic image of FAM-labeled DNA in the presence of unlabeled TCF4 at 0.5 mg/mL, 5% PEG 8000, in buffer S.

**Fig. 8** Proposed TCF4 functioning model. TCF4 typically accumulates in liquid condensates and functions as a trigger for transcription burst when the need arises. Proper TCF4-DNA interaction requires a transient, non-specific interaction between the TCF4 N-terminal IDR and the genome to search for the E-box sequence, and a specific interaction between the TCF4 bHLH domain and the E-box. When DNA binding by TCF4 is impaired due to mutations affecting either DNA-binding residues, stability, or protein localization, TCF4 molecules tend to remain in the condensate, which gradually becomes more solid over time.
