## Supplementary material for "Intrinsically disordered bHLH family member TCF4 drives LLPS in a DNA-dependent manner": figures

**SUPPLEMENTAL FILE**

**Table S1**. Rate constant, *k*, characterizing the dependence of *τ_1/2_* on NaCl concentration.

|  | Incubation time, t | |
| --- | --- | --- |
|  | 3 min | 5 min |
| NaCl (mM) | k (1/s) | |
| 400 | 0.30 ± 0.07 | 0.139 ± 0.016 |
| 500 | 0.19 ± 0.04 | 0.107 ± 0.003 |
| 600 | 0.073 ± 0.018 | 0.045 ± 0.011 |
| 700 | 0.06 ± 0.02 | 0.0447 ± 0.0013 |
| 800 | 0.056 ± 0.018 | 0.023 ± 0.009 |
| 900 | 0.045 ± 0.008 | 0.016 ± 0.004 |

**Table S2**. Rate constant, *k*, characterizing the dependence of *τ_1/2_* on incubation time, *t*.

| NaCl (mM) | | | |
| --- | --- | --- | --- |
| 500 mM | | 750 mM | |
| t (min) | k (1/s) | t (min) | k (1/s) |
| 6.5 | 0.101 ± 0.008 | 6.8 | 0.06 ± 0.02 |
| 8.7 | 0.091 ± 0.019 | 9.2 | 0.0516 ± 0.0012 |
| 11 | 0.107 ± 0.019 | 11.5 | 0.045 ± 0.012 |
| 13.7 | 0.100 ± 0.017 | 14.3 | 0.08 ± 0.02 |
| 17.8 | 0.082 ± 0.014 | 17.8 | 0.054 ± 0.015 |
| 22.5 | 0.09 ± 0.02 | 19.8 | 0.052 ± 0.008 |
| 70 | 0.059 ± 0.004 | 22.3 | 0.038 ± 0.002 |
| 300 | 0.054 ± 0.004 | 24.8 | 0.032 ± 0.008 |
|  |  | 240 | 0.016 ± 0.007 |

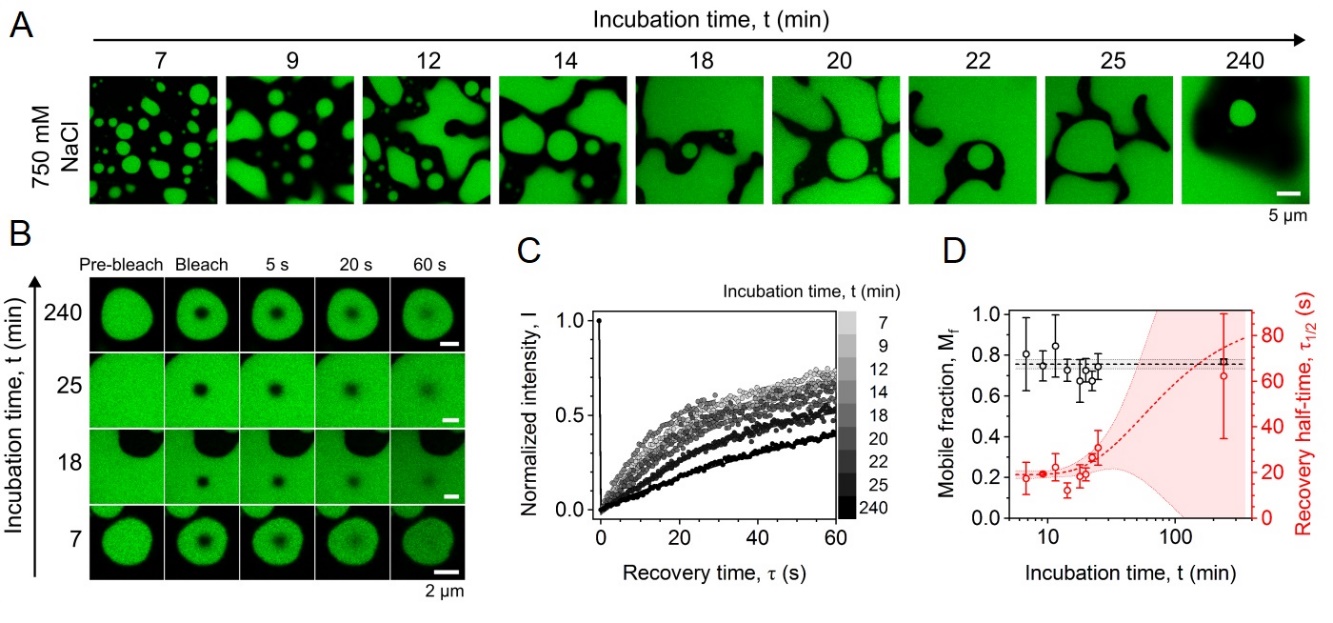

**Fig. S1** (A) Fluorescence confocal microscopy images showing the time evolution of liquid phases formed in solution of TCF4 at 1 mg/mL, at 750 mM NaCl. (B) FRAP images shown for different incubation times. (C) FRAP analysis as a function of incubation time at 750 mM NaCl. (D) Mobile fraction and recovery half-time as a function of incubation time. Data points shown with SD; lines, fitted functions; shaded area, 95% confidence interval. In A and B, the scale bar is the same for all images.

**Table. S3.** Dependency of TCF4 distribution on NaCl concentration using widefield microscopy with DIC. TCF4 was observed as homogenous solution (H), droplets (D) or droplets and aggregates (D+A).

|  | **TCF4 [mg/mL]** | | | |
| --- | --- | --- | --- | --- |
| **NaCl [mM]** | 0.25 | 0.5 | 0.75 | 1.5 |
| 1000 | D+A | D+A | D+A | D+A |
| 900 | D+A | D+A | D+A | D+A |
| 800 | D+A | D+A | D+A | D+A |
| 700 | D | D | D | D |
| 600 | H | D | D | D |
| 500 | H | H | H | H |
| 400 | H | H | H | H |
| 300 | H | H | H | H |
| 200 | H | H | H | H |
| 100 | H | H | H | H |

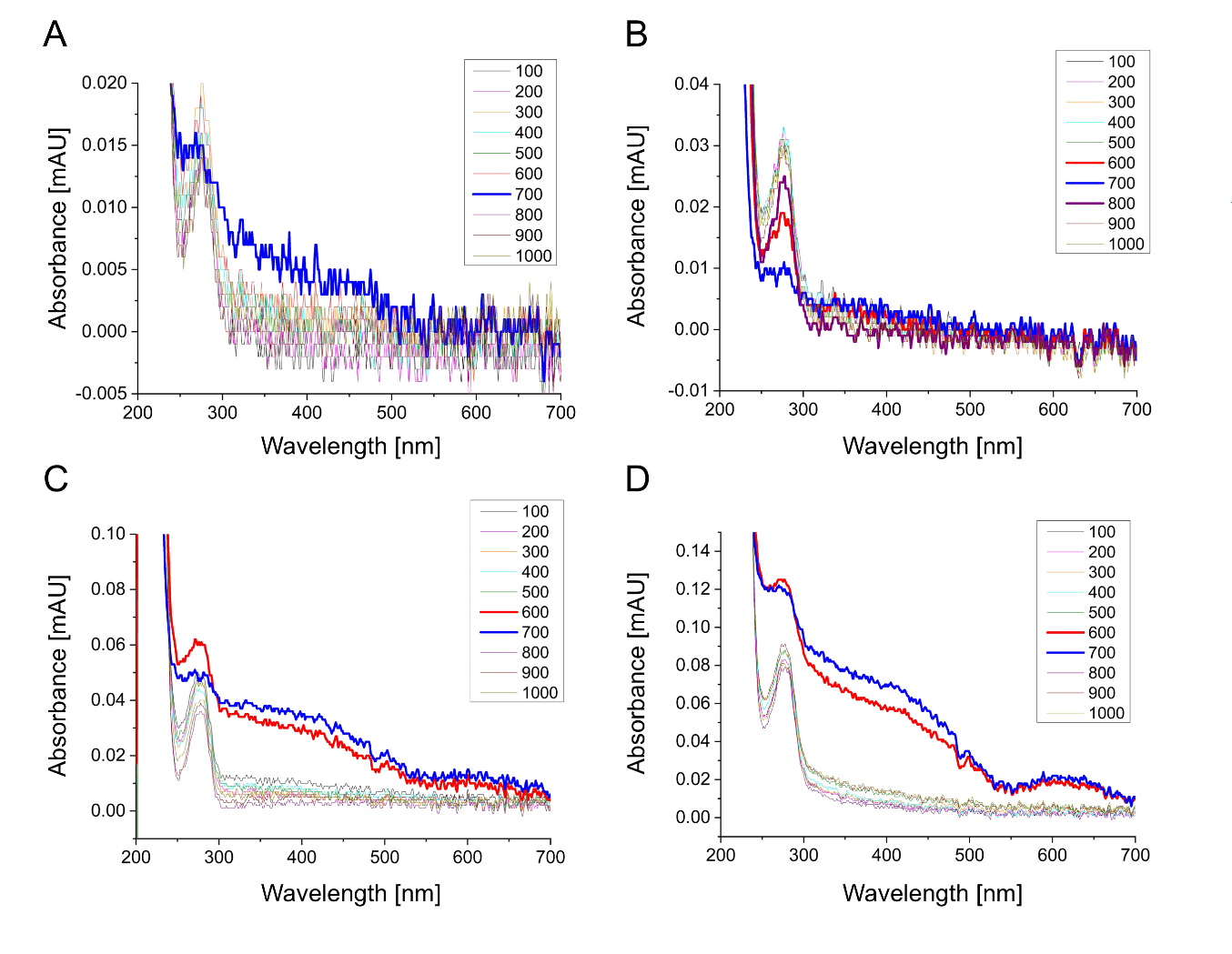
**Fig S2**. TCF4 UV-Vis spectra in different NaCl concentrations. Concentrations of TCF4 were (A) 0.25 mg/mL, (B) 0.5 mg/mL, (C) 0.75 mg/mL, (D) 1.5 mg/mL.

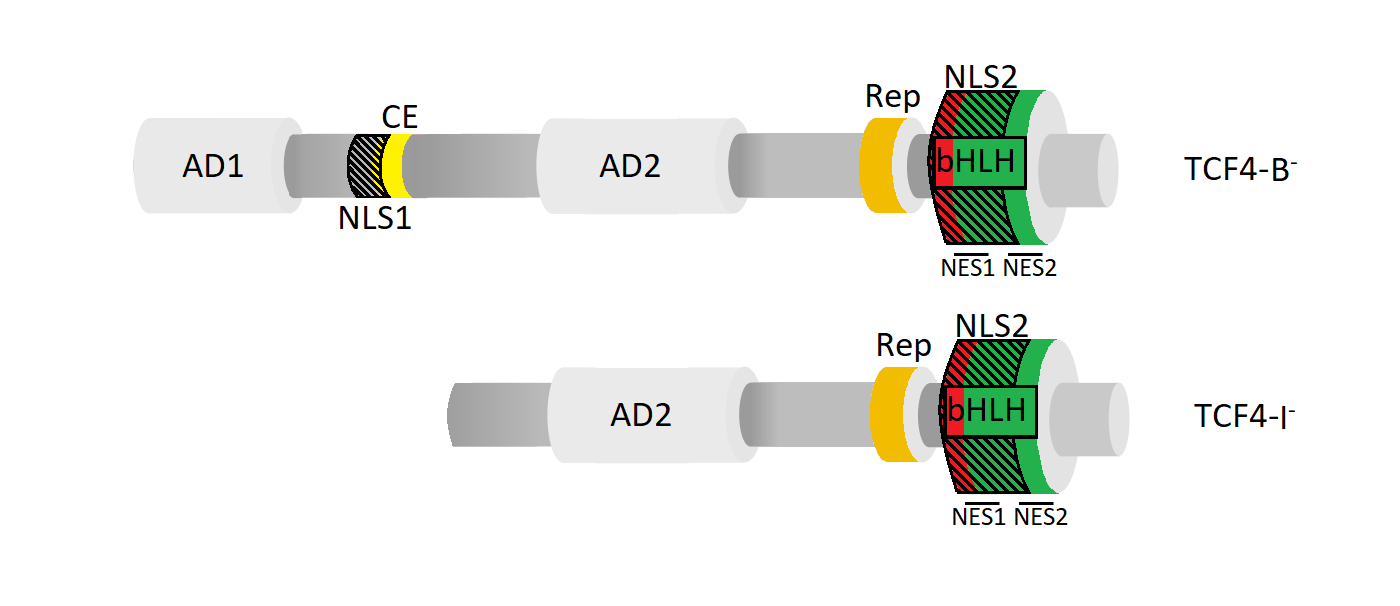

**Fig S3**. Schematic representation of TCF4 canonical isoform TCF4-B ^̶^ , and the shortest human TCF4 isoform, TCF4-I ^̶^. AD1 (1 – 99) and AD2 (325 – 432) are activation domain 1 [1,2] and 2 [1,3], respectively; CE (171 – 185) is conserved element [4] (marked in yellow) ; Rep (511 – 540) is repression domain [5] (marked in blue), bHLH (564 – 617) is basic helix-loop-helix domain [6] (b marked in red and HLH marked in green); NLS1 (156 – 178) and NLS2 (564 – 602) are nuclear localization signal 1 [7] and 2 [8], respectively (black dashed areas); NES1 (585 – 594) and NES2 (600 – 614) are nuclear export signals [8].

1.	Aronheim, A.; Shiran, R.; Rosen, A.; Walker, M.D. The E2A Gene Product Contains Two Separable and Functionally Distinct Transcription Activation Domains. *Proc. Natl. Acad. Sci. U. S. A.* **1993**, doi:10.1073/pnas.90.17.8063.

2.	Massari, M.E.; Jennings, P.A.; Murre, C. The AD1 Transactivation Domain of E2A Contains a Highly Conserved Helix Which Is Required for Its Activity in Both Saccharomyces Cerevisiae and Mammalian Cells. *Mol. Cell. Biol.* **1996**, *16*, 121–129, doi:10.1128/mcb.16.1.121.

3.	Quong, M.W.; Massari, M.E.; Zwart, R.; Murre, C. A New Transcriptional-Activation Motif Restricted to a Class of Helix-Loop-Helix Proteins Is Functionally Conserved in Both Yeast and Mammalian Cells. *Mol. Cell. Biol.* **1993**, *13*, 792–800, doi:10.1128/mcb.13.2.792.

4.	Herbst, A.; Kolligs, F.T. A Conserved Domain in the Transcription Factor ITF-2B Attenuates Its Activity. *Biochem. Biophys. Res. Commun.* **2008**, doi:10.1016/j.bbrc.2008.03.081.

5.	Markus, M.; Du, Z.; Benezra, R. Enhancer-Specific Modulation of E Protein Activity. *J. Biol. Chem.* **2002**, doi:10.1074/jbc.M110659200.

6.	Murre, C.; McCaw, P.S.; Baltimore, D. A New DNA Binding and Dimerization Motif in Immunoglobulin Enhancer Binding, Daughterless, MyoD, and Myc Proteins. *Cell* **1989**, *56*, 777–783, doi:10.1016/0092-8674(89)90682-X.

7.	Sepp, M.; Kannike, K.; Eesmaa, A.; Urb, M. Functional Diversity of Human Basic Helix-Loop-Helix Transcription Factor TCF4 Isoforms Generated by Alternative 5 9 Exon Usage and Splicing. *PLoS One* **2011**, *6*, doi:10.1371/journal.pone.0022138.

8.	Greb-Markiewicz, B.; Kazana, W.; Zarębski, M.; Ożyhar, A. The Subcellular Localization of BHLH Transcription Factor TCF4 Is Mediated by Multiple Nuclear Localization and Nuclear Export Signals. *Sci. Rep.* **2019**, *9*, 15629, doi:10.1038/s41598-019-52239-w.
